# Polypharmacology is an enduring and nearly universal property of kinase inhibitors

**DOI:** 10.64898/2026.03.17.711623

**Authors:** Caitlin E. Mills, Clemens Hug, Karuna Anna Sajeevan, Nicholas Clark, Chiara Victor, Mirra Chung, Sameer Rawat, Hope D’Erasmo, Mariana Pereira Moraes, Bree Aldridge, Mark W. Albers, Ratul Chowdhury, Benjamin M. Gyori, Peter K. Sorger

## Abstract

Despite decades of research, current understanding of the spectrum of targets bound by kinase inhibitors remains incomplete. This complicates mechanism of action studies, drug repurposing, and development of new therapies. Here, we describe kinome-wide profiling of an optimal kinase library (OKL) comprising 192 small molecules selected based on stage of clinical development, chemical diversity, and target coverage. Our results show that polypharmacology is widespread and independent of regulatory approval. The generally understood (“assigned”) targets of approved molecules are not necessarily the most potently inhibited and off-targets include multiple understudied kinases. Moreover, median selectivity has not increased over time We illustrate how an OKL in combination with detailed kinome profiling can be used to identify potential toxicity targets, repurpose anti-inflammatory drugs for neurodegenerative and infectious diseases, and perform chemical genetic studies. Our studies also highlight how much remains to be discovered about the chemistry and biology of one of the largest classes of human therapeutics.

## INTRODUCTION

Optimizing the selective binding of a chemical compound to a desired protein target is a key step in contemporary small molecule drug development. This is particularly true in the case of targets that are members of multi-protein families. The human kinome comprises ∼530 proteins (with the number somewhat dependent on definition)^1^ and as of 2026, over 100 kinase inhibitors have been FDA approved for use as therapeutics^2^ with an additional 369 compounds against 129 targets undergoing clinical trials^3,4^. This represents one of the most active areas of drug discovery and there were more kinase inhibitors approved by the FDA in 2025 than in any prior year^5^. In current practice, kinase inhibitors are most commonly designed to bind one or a small set of closely related members of the kinome (we refer to these as “assigned” targets). It is therefore common to describe drug ‘*A*’ as an inhibitor of assigned target ‘*B*’, but it is generally understood that binding to other targets (polypharmacology) plays an essential – if poorly understood – role in drug activity. Although kinase inhibitors (like all therapeutics) achieve regulatory approval based on clinical endpoints, survival in the case of anti-cancer drugs, a description of mechanism of action is an essential component of regulatory filings, and this invariably emphasizes the presumed role of assigned target(s). A complete understanding of how inhibition of other potential targets contributes to the clinical success of small molecules often occurs post-approval and is broadly dependent on better ways of assessing polypharmacology.

Both academic and commercial platforms exist to profile the binding affinities of small molecules against the great majority (∼75%) of all protein kinases. Several high-impact projects have studied the polypharmacology of kinase inhibitors at scale^6–10^ using compound libraries that vary in composition. However, despite a steady stream of drug approvals, it has been some time since kinase inhibitor polypharmacology has been studied in depth. Moreover, new commercial resources and cheminformatic tools now make it easier to assemble libraries that strike an optimized balance between chemical diversity, selectivity, stage of clinical development, and library size (and thus, cost)^11^. The 192 compound Optimal Kinase Library (OKL) in this paper is one example of a library optimized on several competing characteristics.

The literature on protein kinase inhibitors is large, and affinity data obtained from the literature are actively curated in public databases such as ChEMBL^12^, IDG Pharos^13^ and BindingDB^14^. Metrics such as affinity spectra (TAS)^11^ consolidate available data on inhibitor binding and non-binding relationships in public data while accounting for uncertainty in the underlying evidence. However, systematic analysis of polypharmacology is not as simple as assembling literature knowledge. For kinase inhibitors, this literature relies on a wide range of assay types (e.g. enzymatic inhibition, affinity mass spectrometry, binding to recombinant protein libraries) and conditions (e.g. ATP or substrate concentrations) with few overlapping data points. This makes kinome-wide analysis difficult. Moreover, affinity screening is often biased toward assigned target(s), suspected off-targets, and “toxicity” targets known to limit clinical utility. Data on “known non-binders” is particularly sparse, even though it is essential for establishing selectivity^11^. Thus, existing knowledge is too incomplete to answer basic questions about the breadth and significance of compound polypharmacology.

In this paper we combine an OKL with the most comprehensive and widely used kinome profiling technologies (Eurofins’ KINOMEscan platform)^15^ to collect kinome-wide affinity data at four concentrations for 192 kinase inhibitors optimized for selectivity, affinity, and approval status. We introduce a Bayesian inference framework to estimate K_d_s for all inhibitor-kinase pairs, enabling inference of affinities outside tested concentrations. We find that OKL compounds inhibit all assayed kinases at 1 µM or below (and 94% coverage 100 nM or below) but with widely varying various degrees of polypharmacology. For many therapeutic drugs, the assigned target was not the only, or even the highest affinity target. Kinase clustering based on measured polypharmacology differs from the familiar view of the kinome tree based on sequence similarity, demonstrating that sequence alignment is an imperfect way of thinking about target affinity. We illustrate the utility of the fully annotated OKL as a screening tool by using it to dissect, identify and characterize neuroprotective agents active in a model of Alzheimer’s disease, investigate vulnerabilities in a model of cisplatin-resistant ovarian cancer, and identify inhibitors of non-mammalian pathogenic kinases. These use cases show that systematic kinome-scale knowledge of kinase polypharmacology has broad utility across biological contexts.

## RESULTS

### Establishment of kinome-wide affinities for a practically scaled Optimal Kinase Library

The larger and more diverse a chemical library and the more comprehensive the data on activity the greater the insight into compound-target relationships. However, collecting kinome-wide affinity data is expensive; for 10-point enzymatic assays this is ∼$100/curve or $10M for 200 compounds against 500 targets. The KINOMEscan® kinome-scale profiling assay available from Eurofins is 5-10-fold cheaper and has been used in more than 2,000 publications. Systematic profiling is nonetheless sufficiently expensive to require compact libraries^6–9,11,16–18^. Thus, the OKL in this work aimed for a minimal library balancing target coverage and structural diversity so that, ideally, each kinase was bound by two inhibitors with the highest possible selectivity and the greatest difference in chemical structure (i.e. Tanimoto similarity ≤ 0.2 between pairs based on Morgan fingerprints with a two bond radius)^11^. When a choice was available, we selected compounds that were approved for use in humans. This resulted in a 192-compound library comprising 44 FDA-approved drugs, 98 molecules that have been, or are currently, in clinical trials, and 50 pre-clinical “tool” compounds having a total of 115 unique assigned targets (**Figure 1a-b, Supplementary Table 1).** Overlap between libraries previously subjected to systematic kinome profiling and OKL was surprisingly limited (**Figure 1c**) due to differences in study goals, our emphasis on target and chemical diversity, and a steady increase in the scope of clinical approvals^6–9,17^. Overlap in the kinases profiled also varied due the types of assays run and advances in methods. For these reasons, and those outlined above, knowledge of kinome-wide binding by OKL inhibitors was sparse (**Figure 1d**).

**Figure 1:**
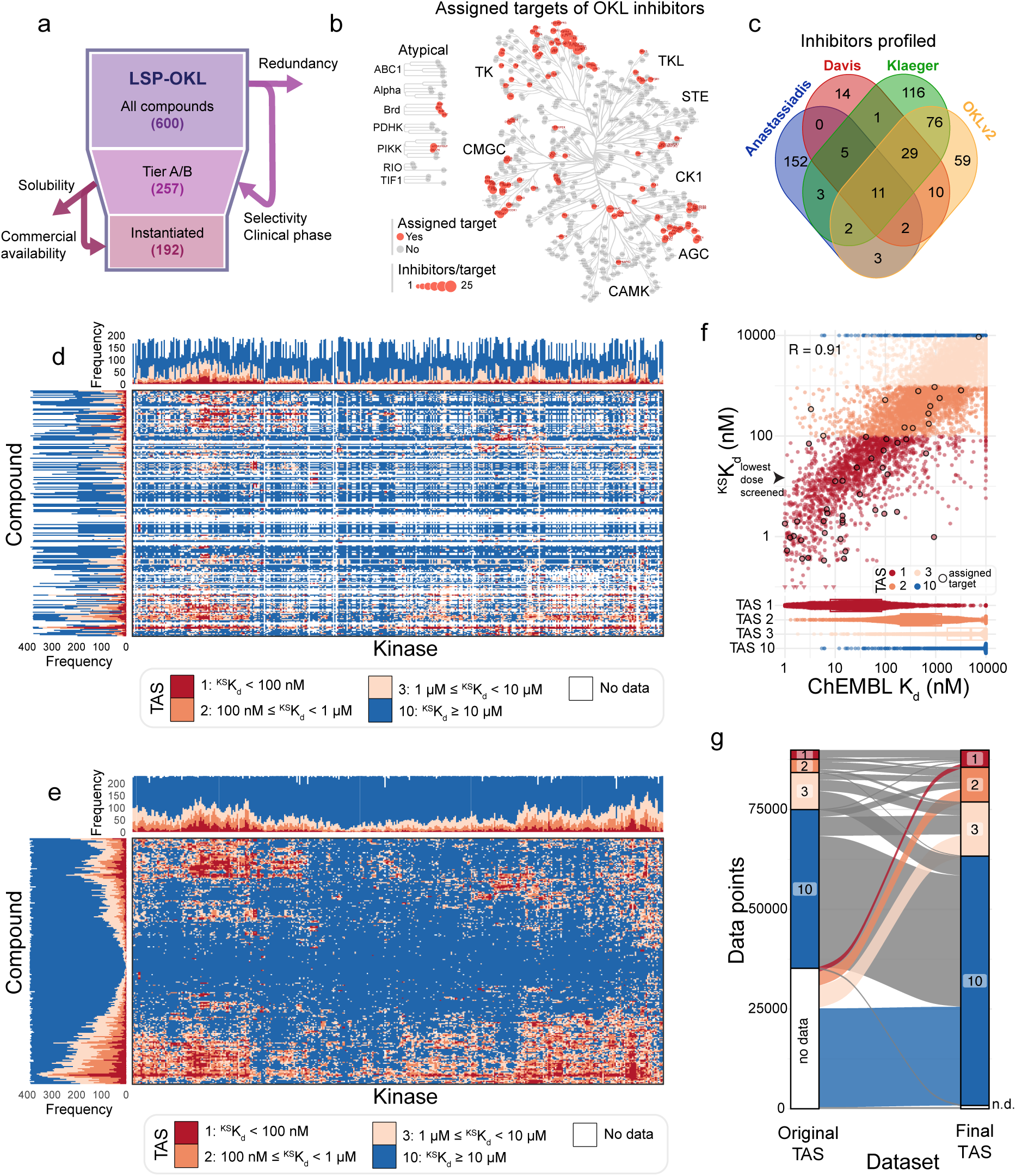
A complete view of kinome-wide affinities for the 192 compound ‘Optimal Kinase Library’. **(a)** Schematic representation of the reduction of OKL from 600 potential compounds to an instantiated OKL set of 192 compounds. **(b)** The assigned targets of OKL compounds shown on the kinome tree; red indicates an assigned target and the size of the spot indicates the number of inhibitors per target. **(c)** Venn diagram showing the overlap between the inhibitors profiled in previous kinome-wide profiling projects and those in this paper. **(d)** Kinome-wide target affinity spectra based on ChEMBL data for all OKL compounds prior to the current study. **(e)** Kinome-wide target affinity spectra for all OKL compounds after the addition of the data generated in the current study. **(f)** Scatterplot comparing the ^KS^K_d_ values measured in this paper to those previously available in ChEMBL. Data points are colored by TAS value, and the values for each compound’s assigned target(s) are outlined in black. **(g)** An alluvial plot depicting the knowledge gained from addition of the data generated in the current study. TAS values on the left are from (c) and on the right from (d).

KINOMEscan scanMAX profiling (hereafter KINOMEscan) is based on a competitive binding assay comprising 406 wildtype (WT) human kinases and 59 kinases carrying oncogenic or resistance mutations^19,20^ plus three non-mammalian kinases (the availability of kinases is dependent in part on the feasibility of making them recombinantly; **Supplementary Table 2**). In KINOMEscan, a compound of interest is incubated with oligo-tagged recombinant kinases and immobilized ATP-like ligands capable of binding all kinases in the library; the amount of ligand-bound kinase is measured by qPCR to generate a ‘percent of control’ value (relative to incubation with DMSO); a value of 0 denotes strong binding and 100 no binding. We assayed each OKL inhibitor at four concentrations spanning four orders of magnitude (10 µM, 1 µM, 100 nM and 12.5 nM) to generate 89,856 four-point dose response curves (192 OKL compounds x 468 targets). Quality control (see Methods, **Supplementary Figure 1a-g**) revealed ∼900 *discordant* dose response profiles (∼1% of the data) in which percent control did not fall with increasing inhibitor concentration (**Supplementary Table 3**). These data were omitted from further analysis.

We explored a variety of approaches for estimating KINOMEscan-derived dissociation constants (^KS^K_d_) and settled on Bayesian inference. Bayesian modeling substantially improved the reliability of (^KS^K_d_) estimations compared to linear interpolation, particularly when values were below the lowest concentration tested experimentally (12.5 nM; **Supplementary Figure 1h,** see **Supplementary Note**) and resulted in a more complete view of kinome-wide binding for OKL inhibitors (**Figure 1e, Supplementary Table 4**). Those inhibitor-kinase pairs for which K_d_ data from biochemical assays were available in ChEMBL exhibited good agreement (R = 0.91) with ^KS^K_d_ estimates (**Figure 1f-g**), providing independent confirmation of OKL-KINOMEscan values. In contrast, the most similar existing dataset (with respect to library composition; labelled Klaeger in **Figure 1c**)^6^ exhibited a correlation with ChEMBL data of R=0.64. Two key features of a library of small molecule inhibitors are coverage – how many targets are bound by how many inhibitors; and selectivity – how many targets are preferentially (selectively) bound by any inhibitor. OKL achieve full coverage of all 468 assayed targets at a relatively high inhibitor concentration of 1 µM or below (2-130 inhibitors per kinase at ^KS^K_d_ ≤ 1 µM) and 94% coverage of WT kinases at 100 nM or below (0-57 inhibitors for 379/406 WT kinases at ^KS^K_d_ ≤ 100 nM); at this lower concentration all mutant kinases were also inhibited, likely because they have been the focus of years of intensive medicinal chemistry (**Figure 2a-b, Supplementary Figure 2a-c, Supplementary Figure 3**).

**Figure 2:**
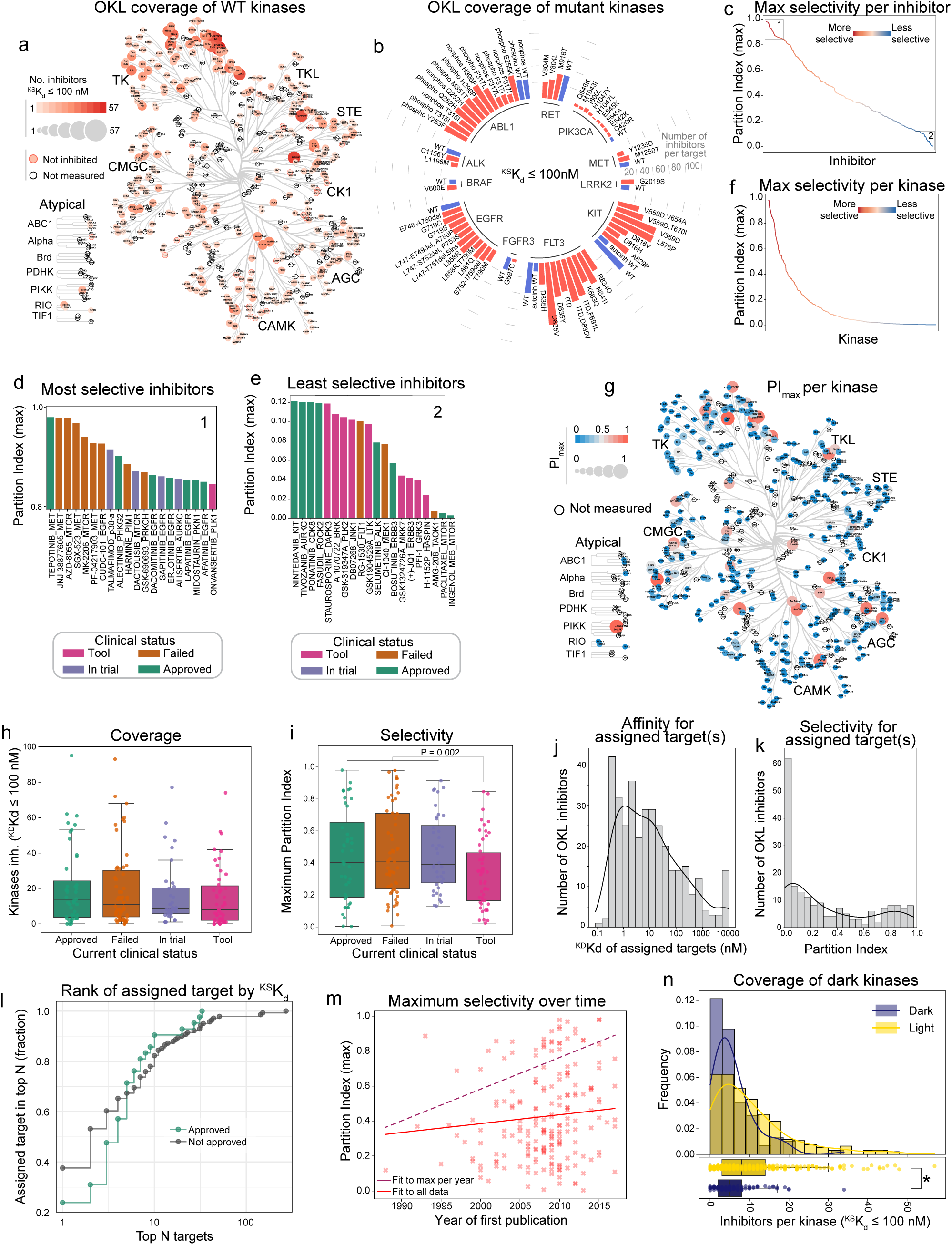
Kinome coverage and selectivity of OKL inhibitors. **(a)** Number of OKL inhibitors (^KS^K_d_ < 100 nM) per kinase shown on the kinome tree by the size and intensity of the markers. Kinases not inhibited are outlined in red, kinases not assayed in the KINOMEscan panel are outlined in black. **(b)** Radial bar plot showing the number of OKL inhibitors with ^KS^K_d_ ≤ 100 nM for all mutant kinases (red bars) assayed in the KINOMEscan panel. WT kinases are denoted by blue bars. **(c)** The maximum partition index (PI_max_) for each inhibitor in OKL. **(d)** Barplot of the most (box 1 in panel c) and **(e)** least (box 2 in panel c) selective inhibitors and associated kinase target colored by current clinical status. **(f)** PI_max_ for each kinase. **(g)** CORAL tree depicting the PI_max_ for each kinase, the larger and redder the circle, the more selectively that kinase is inhibited by an OKL compound. **(h)** Boxplot showing the number of kinases per inhibitor with ^KS^K_d_ ≤ 100 nM by clinical status. **(i)** Boxplot showing the maximum partition index for each OKL inhibitor by clinical status. P-value is from a Welch’s t-test comparing tool compounds to all others. **(j)** Histogram of the ^KS^K_d_ values for the assigned target(s) of OKL compounds. **(k)** Histogram of the partition indices for the assigned target(s) of OKL compounds. The PI values were added together for inhibitors with multiple assigned targets. **(l)** Cumulative stairstep plot showing how often the nominal target is in the top N targets by ^KS^K_d_ for approved (green) and not approved (gray) inhibitors. **(m)** Scatterplot showing PI_max_ with respect to the year each compound was first published. The lines of best fit to all data points and to the highest datapoint per year are shown in solid red and dashed maroon, respectively. **(n)** Histogram showing the number of OKL inhibitors that bind each dark and illuminated kinase with ^KS^K_d_ ≤ 100 nM. Boxplots representing the same data are shown below the histograms. The * denotes Welch’s t-test P < 0.0001.

### Kinase inhibitor selectivity for tool compounds and human therapeutics

Several metrics are available for quantifying library selectivity^6,21,22^; we used the easily-understood partition index (PI), which represents the fraction of an inhibitor bound to each kinase (for which a ^KS^K_d_ estimate was available) in a theoretical situation in which all kinases are present in excess^21^. A perfectly selective inhibitor binding a single target would have a maximum PI (PI_max_) = 1 and an inhibitor binding many targets with similar affinity would have PI_max_ ≪ 1. PI_max_ values for OKL inhibitors spanned a range from < 0.1 for AMG-208 (whose development was abandoned after phase 1) to > 0.98 for tepotinib (a therapeutic for non-small cell lung cancer that binds the MET receptor tyrosine kinase; RTK) (**Figure 2c-e**). We identified 69 “highly selective” compounds that bound to 37 WT kinases with PI_max_ > 0.5, but maximum selectivity for most kinases was PI_max_ < 0.2 (**Figure 2f-g; Supplementary Table 5**). Overall, we found that polypharmacology was widespread among human therapeutics. With respect to use of kinase inhibitors in research, it has been proposed that reliable results can be obtained only if molecules meet the selectivity definition of a “chemical probe”^23^. Sixteen compounds in the OKL collection meet this definition (^KS^K_d_ < 100 nM, and CATDS_most-potent_ > 0.5 for a single target; **Supplementary Table 6**).

Using OKL-KINOMEscan data, we examined four prevailing assumptions about the development of kinase inhibitors as human therapeutics: (i) approved drugs are more selective than compounds that failed in clinical trials (where ‘failed’ is defined as a compound tested at least through phase 1 but neither approved nor currently under clinical investigation^3^); (ii) assigned targets are generally potently and selectively inhibited, particularly for FDA-approved drugs; (iii) compound selectivity has increased over time; and (iv) understudied (dark) kinases (per the definition of the NIH Investigating the Druggable Genome project)^24^ are less widely inhibited than well-characterized (illuminated) kinases.

First, we grouped compounds based on maximum reported clinical trial phase, approval status, or trials registered in ClinicalTrials.gov and found that FDA-approved drugs do not differ from compounds in other stages of clinical development in terms of coverage or selectivity; however, purely pre-clinical tool compounds are on average less selective (**Figure 2h-i, Supplementary Figure 2d-e**). Second, while about half of assigned targets are bound with high affinity (**Figure 2g**, ^KS^K_d_ ≤ 10 nM for 48% of assigned targets), selectivity varies widely (PI = 0 to 0.98, median 0.06, mean 0.17; **Figure 2k**) suggesting that many compounds are potent but not highly selective. Considering only FDA-approved drugs, an assigned target was the *highest* affinity target in only 23% of cases examined; for compounds that are not FDA-approved the proportion was 37% (**Figure 2l**). Thus, if we define a promiscuous inhibitor as one that that binds many kinases, and a promiscuous kinase as one that binds many inhibitors, profiling of the OKL library revealed a wide range of promiscuity in inhibitors and kinases, even among FDA-approved drugs. Moreover, if we use the year of first publication to estimate the “age” of an OKL compound we find that average selectivity has not changed significantly over time. Thus, high selectivity (PI_max_ value) has not been necessary for clinical success with kinase inhibitors. However, the highest selectivity achieved by a single compound has increased over time (**Figure 2m**; dark dotted line) presumably arising from advances in kinase inhibitor medicinal chemistry.

### Kinase anti-targets and dark kinases

The concept of an anti-target, a protein that must be spared during drug discovery, is exemplified by the cardiac K+ channel hERG^25^. It has been proposed that some kinases also represent anti-targets,^26,27^ although no definitive list has as-yet been assembled. One striking and unreported example of a potential anti-target interaction in OKL-KINOMEscan data involves alectinib, which is approved as Alecensa® for metastatic non-small cell lung cancer and has ALK as an assigned target. OKL-KINOMEscan suggests that alectinib can also bind PHKG2, the liver-specific γ-subunit of the phosphorylase kinase. Enzymatic assays confirmed that alectinib is an IC50 = 33 nM inhibitor of PHKG2 (IC50 = 0.64 nM for ALK; C_max_ in patients is 1.38 µM^28^; **Supplementary Figure 2f**). Mutation in PHKG2 give rise to severe liver disease^29^ and its inhibition by sunitinib causes liver toxicity^30^. Hepatotoxicity is a recognized problem in up to 60% of patients treated with alectinib^31^ (with grade 2-4 toxicity in 5-8% of patients) and we speculate that this is a consequence of inhibition of PHKG2. Examination of ^KS^K_d_ values for ALK inhibitors across the OKL-KINOMEscan dataset shows that whereas as some target PHKG2 others do not. Thus, it should therefore be possible to create potent ALK inhibitors in which PHKG2 is not bound, potentially reducing hepatoxicity.

Wide differences in the levels of attention given to kinases over time has given rise to the concept of “understudied” or dark kinases^32^ (the focus of a major NIH program)^33^. We found that these understudied kinases^24^ were widely bound and selectively inhibited even by FDA-approved compounds (**Figure 2n, Supplementary Figure 2g, h**). The number of OKL compounds inhibiting dark kinases was less than for illuminated kinases however (median of 5 vs 8 inhibitors per kinase at a threshold of ^KS^K_d_ ≤ 100 nM, respectively). These largely uncharacterized inhibitory activities represent a potential starting point for targeting understudied kinases and may also provide insight into efficacious off-target activities of existing human therapeutics.

### Kinase promiscuity is associated with a DFG-out conformation binding Type I and II inhibitors

What is the basis of promiscuous binding of some human kinases to OKL inhibitors? To address this, we focused on the 18 most promiscuous kinases in our dataset (15 TKs, two STE kinases, and one CAM kinase; **Figure 3a**) each of which bound ≥ 33 compounds with relatively high affinity (^KS^K_d_ ≤ 100 nM). Overlap among these inhibitors was limited with two or fewer of the same compounds binding to 16/18 promiscuous kinases (**Figure 3b**, blue points); for example, MEK5 (**Figure 3b**, red point) was bound by five inhibitors that did not bind any other promiscuous kinases (at ^KS^K_d_ ≤ 100 nM). Thus, promiscuous kinases do not simply bind a common set of promiscuous inhibitors.

**Figure 3:**
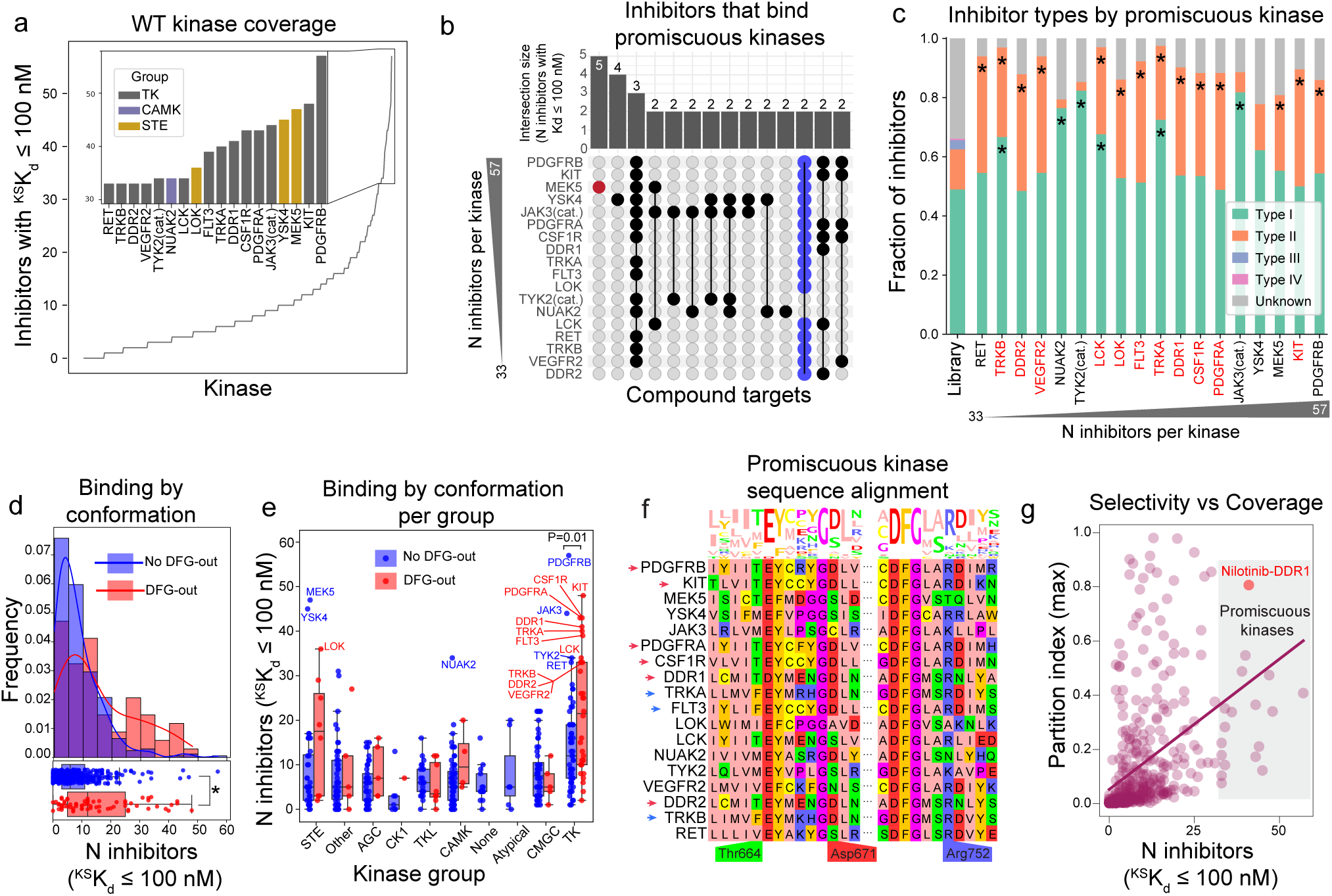
A subset of kinases are promiscuous for OKL inhibitors. **(a)** The number of OKL inhibitors with ^KS^K_d_ ≤ 100 nM per kinase. The top 15 ranks with ties are shown in the inset bar plot colored by kinase group. These 18 kinases are referred to as promiscuous. **(b)** Upset plot showing the number of overlapping inhibitors with ^KS^K_d_ ≤ 100 nM for the promiscuous kinases. **(c)** Stacked bar graph showing the distribution of inhibitor types in OKL and that bind each promiscuous kinase. Kinases identified as promiscuous in a previous study are in red. **(d)** Histograms of the number of inhibitors per kinase with ^KS^K_d_ ≤ 100 nM for kinases with (red) and without (blue) reported DFG-out structures. KDE plots are overlaid for visualization. Boxplots representing the same data are shown below the histograms. The * denotes Welch’s t-test P < 0.0001. **(e)** Boxplots of the number of inhibitors with ^KS^K_d_ ≤ 100 nM for kinases having (red) and not having (blue) a DFG-out structure in the PDB by kinase group. Welch’s t-tests were performed on all pairs, P-values are only shown where P < 0.05. **(f)** Multiple sequence alignment and sequence logo plot of the promiscuous kinases. Residues previously implicated in RTK promiscuity are labeled. Amino acids are colored based on the zappo color scheme. **(g)** Scatter plot of the number of inhibitors per kinase with ^KS^K_d_ ≤ 100 nM with respect to the kinase’s maximum partition index. Promiscuous kinases are those found with the region donated by a gray box.

Identified structural determinants of binding specificity for ATP-competitive kinase inhibitors involve the conformation and local environment of a conserved three amino acid motif (DFG) that lies immediately N-terminal to the activation loop^34^. Kinases assume both a *DFG-in* conformation, which is generally catalytically active, and a *DFG-out* conformation, which is inactive. Kinase inhibitors are classified into type I inhibitors that bind both DFG-in and DFG-out conformations and type II inhibitors that bind only DFG-out conformations; type III and IV inhibitors (so-called “allosteric inhibitors”) bind outside the active site and are substantially less common^35^. We found that 14/18 promiscuous kinases were bound by more type II inhibitors than expected by chance (given OKL composition) (**Figure 3c**) consistent with binding to a DFG-out conformation. Moreover, kinases with a DFG-out structure in the Protein Data Bank (PDB)^36^ were more promiscuous than those without (**Figure 3d**, mean number of ^KD^K_d_ ≤ 100 nM inhibitors 15.5 and 8.3, respectively; Welch’s t-test P = 2.66 x 10^-5^). This difference was greatest for kinases in the STE and TK families (**Figure 3e**).

Six RTKs promiscuous for OKL compounds (PDGFRA, PDGFRB, KIT, CSF1R, DDR1, DDR2) were previously shown to bind many members of the 645-compound PKIS2 library, which has no overlap with OKL^37,16^. This was attributed to stabilization of the DFG-out conformation by a D671 to R752 salt bridge (the numbering is based on the DDR1 kinase)^37^ (**Figure 3f**, red arrows). In our study, three additional RTKs (TRKA, TRKB and FLT3) having analogous D671 and R752 residues (**Figure 3f**, blue arrows) were also promiscuous. However, the same residue pair is present in four kinases that are not RTKs and are not promiscuous, suggesting the impact of the D671 to R752 salt bridge on inhibitor binding is limited to RTKs (**Supplementary Table 7**). Since several of these proteins are being targeted for development of human therapeutics, we asked whether promiscuity is incompatible with selective inhibition. Our data suggest that the answer is no: DDR1, for example, is bound by 41 OKL compounds (^KS^K_d_ ≤ 100 nM) but one of these (Nilotinib, whose assigned target is BCR-ABL) is highly selective, with a partition index of 0.81 (**Figure 3g**, red point).

### Sequence homology and inhibitor binding provide distinct views of the kinome

The classic kinome tree is based on a classification scheme popularized by Manning, et al.^38^ in which kinases are clustered based on sequence homology (primarily in the catalytic domain). The tree is split into nine major groups (e.g. TKs, CAM kinases, AGC kinases, etc.) that each contain multiple families. Targeting specific isoforms within these families is a common goal of drug discovery, as exemplified by the search for PI3K inhibitors that discriminate between oncogenic kinases and immune regulators^39^. To see if OKL affinity data could assist in similar tasks, we performed hierarchical clustering based on the Spearman correlation of ^KS^K_d_ values. This yielded nine major clusters (**Figure 4a, b**) that differed substantially from the groups generated by sequence alignment (**Figure 4b**). Taking the CMGC group as an example (**Figure 4b**, dotted outline), the 14 kinases in the MAPK family form one sequence-based cluster but six OKL clusters (**Figure 4b**, blue branch; **Figure 4c**). Thus, proteins most related in sequence can have very different inhibitor binding profiles: ERK1 and ERK2 are bound by different OKL inhibitors than ERK3 and ERK4, for example. In contrast, a second branch in the CMGC group comprising DYRK and SRPK family kinases lie in a single OKL cluster with a single outlier: HIPK4 (**Figure 4b**, red branch; **Figure 4d**). A HIPK4 inhibitor (**Figure 4e**) is more likely to bind other kinases in OKL cluster 3 which includes c-Jun N-terminal kinases (JNK1, JNK2, JNK3) which are in the MAPK family (**Figure 4a-d**). Inhibitor selectivity therefore represents a different way of viewing the kinome tree than sequence similarity and can be used to prioritize off-targets for deeper analysis. With additional data, it should be possible to extend this idea and construct a kinome tree based entirely on small molecule binding as opposed to sequence similarity.

**Figure 4:**
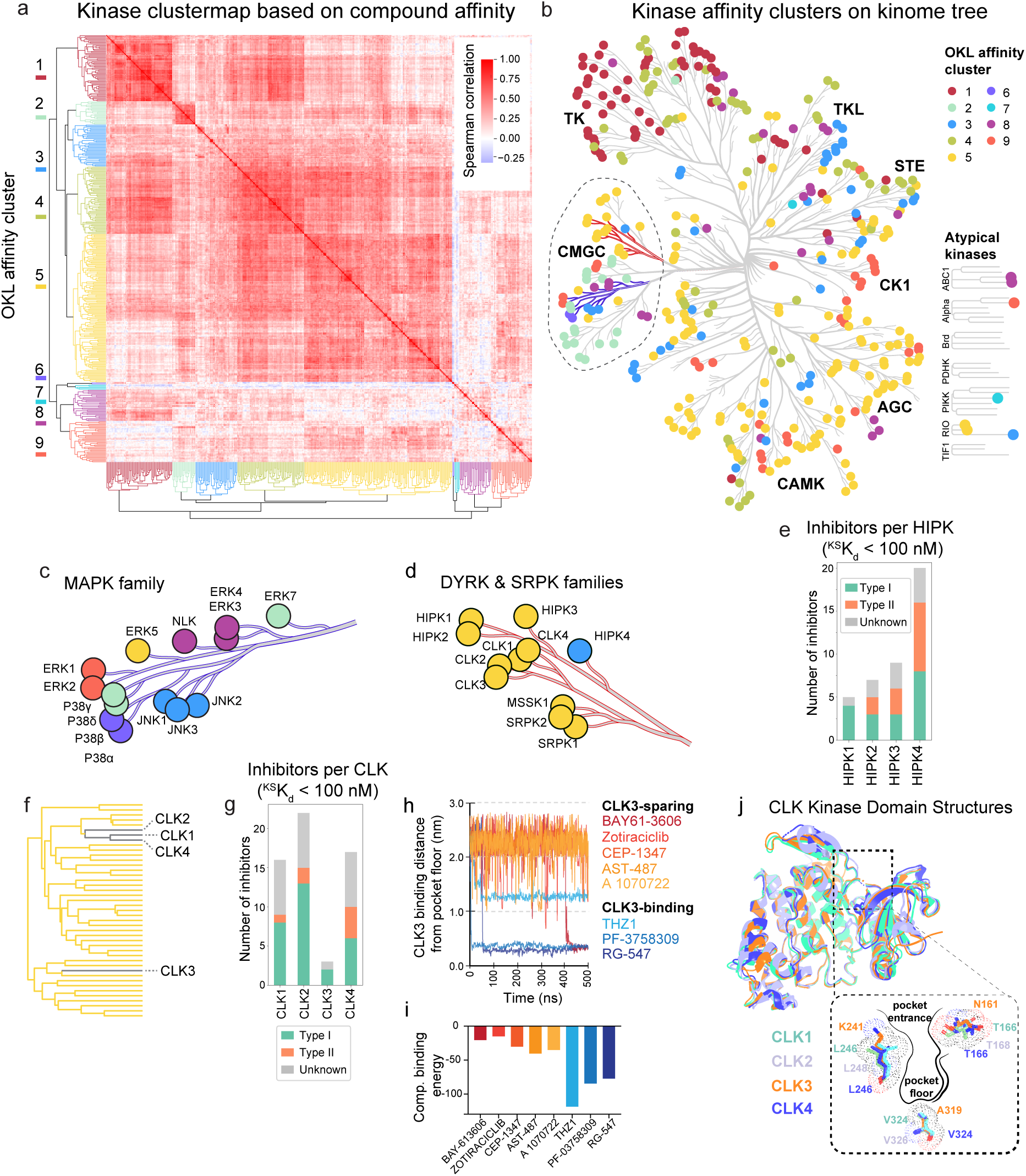
Kinase sequence homology with respect to OKL compound affinities. **(a)** Hierarchical clustermap of the Spearman correlation between kinases based on their ^KS^K_d_ values for all OKL compounds. **(b)** The colors of the clusters are overlaid on the kinome tree. The blue branches are magnified in **(c)** and the red branches in **(d)**. **(e)** Bar plot of the number of OKL inhibitors with ^KS^K_d_ ≤ 100 nM for each HIPK colored by inhibitor type. **(f)** Subset of the hierarchical tree from (a) showing the positions of each CLK. **(g)** Bar plot of the number of OKL inhibitors with ^KS^K_d_ ≤ 100 nM for each CLK colored by inhibitor type. **(h)** 500ns molecular dynamics trajectories showing the distance of the indicated inhibitors from the floor of CLK3’s binding pocket. **(i)** Computational binding energy scores from GROMACS molecular dynamics simulations of eight inhibitors with CLK3. **(j)** A visual overlay of the overall structures of CLK1,2,3, and 4 and zoomed-in view of their binding pockets highlighting key amino acids.

The CDC2-like kinases (CLKs) have been linked to diseases ranging from cancer to neurodegeneration^40^, and CLK inhibitors are under investigation for multiple types of cancer. CLK1, CLK2, and CLK4 cluster tightly together based on OKL binding profiles but CLK3 - like HIPK4 - is an outlier relative to its closest homologues (**Figure 4f, g**). To investigate residues involved in small molecule binding by CLK3, we performed 500 ns all-atom molecular dynamics (MD) simulations for eight inhibitors having a range of affinities for CLK kinases. Starting the simulation with compounds outside the binding pocket, we found that high affinity CLK3 binders entered and remained in the pocket as measured by the distance from residue A319 on the pocket floor (< 0.4 nm for PF-3758309 and RG-547, and ∼1.2 nm for THZ1; **Figure 4h**) and this corresponded to computational binding scores < −75 (**Figure 4i**). In contrast, inhibitors with ^KD^K_d_ ≥ 1000 nM for CLK3 remained distant (∼2.5 nm) from the pocket floor over the 500 ns simulation suggesting that they failed to enter the pocket, thereby reproducing the experimental data. Simulations suggested that two residues differing between CLK3 and other CLKs at the pocket entrance are likely involved in this selectivity: T166→N161, L246→K241 (number based on CLK3 residue positions; **Figure 4j**). These residue positions are known, along with V324→A319 on the pocket floor, to affect the charge distribution and size of the drug/ATP binding pocket^41,42^. Thus, subtle structural differences in kinase binding pockets that are not discernable by sequence alignment can be detected by OKL kinome profiling and potentially used to guide isoform-selective inhibitor development.

### Using an optimized kinase library and affinity data to explore neurodegeneration

To explore practical uses of OKL data, we performed cell-based screening in three distinct biological contexts: (i) rescue of neuronal cell death in a model of Alzheimer’s disease (ii) identifying targets for killing of platinum resistant ovarian cancer cells and (iii) identification of inhibitors of non-mammalian kinases. A subset of AD is thought to involve neuronal death triggered by an inflammatory response to cytoplasmic double stranded RNA (cdsRNA)^43^. This can be recapitulated in a cell-based screen by treating differentiated ReNVM neurons with the dsRNA mimetic, polyI:C, which leads to type I interferon signaling and cell death (**Figure 5a**). Neuronal death can be rescued to variable degrees by Janus family kinase inhibitors (the JAK family comprises JAK1-3 and TYK2)^43,44^. To determine if off-targets are involved in rescue, we extended the OKL dataset by performing KINOMEscan profiling on six additional clinically advanced JAK inhibitors, focusing on the two most informative concentrations: 100 nM and 10 µM (**Supplementary Table 8**); the resulting OKL+JAKi dataset included a total of 21 inhibitors with assigned or reported targets in the JAK family. We then screened the OKL at 1 µM in ReNVM cells treated with polyI:C and identified 14 inhibitors that conferred a significant (20% or greater) increase in viability (**Figure 5b**). In the subsequent rescreen, these 14 hits, and all compounds with one or more assigned targets in the JAK family (33 compounds in total), were then tested for rescue at four doses (10, 1, 0.1, 0.01 µM) (**Figure 5c**).

**Figure 5:**
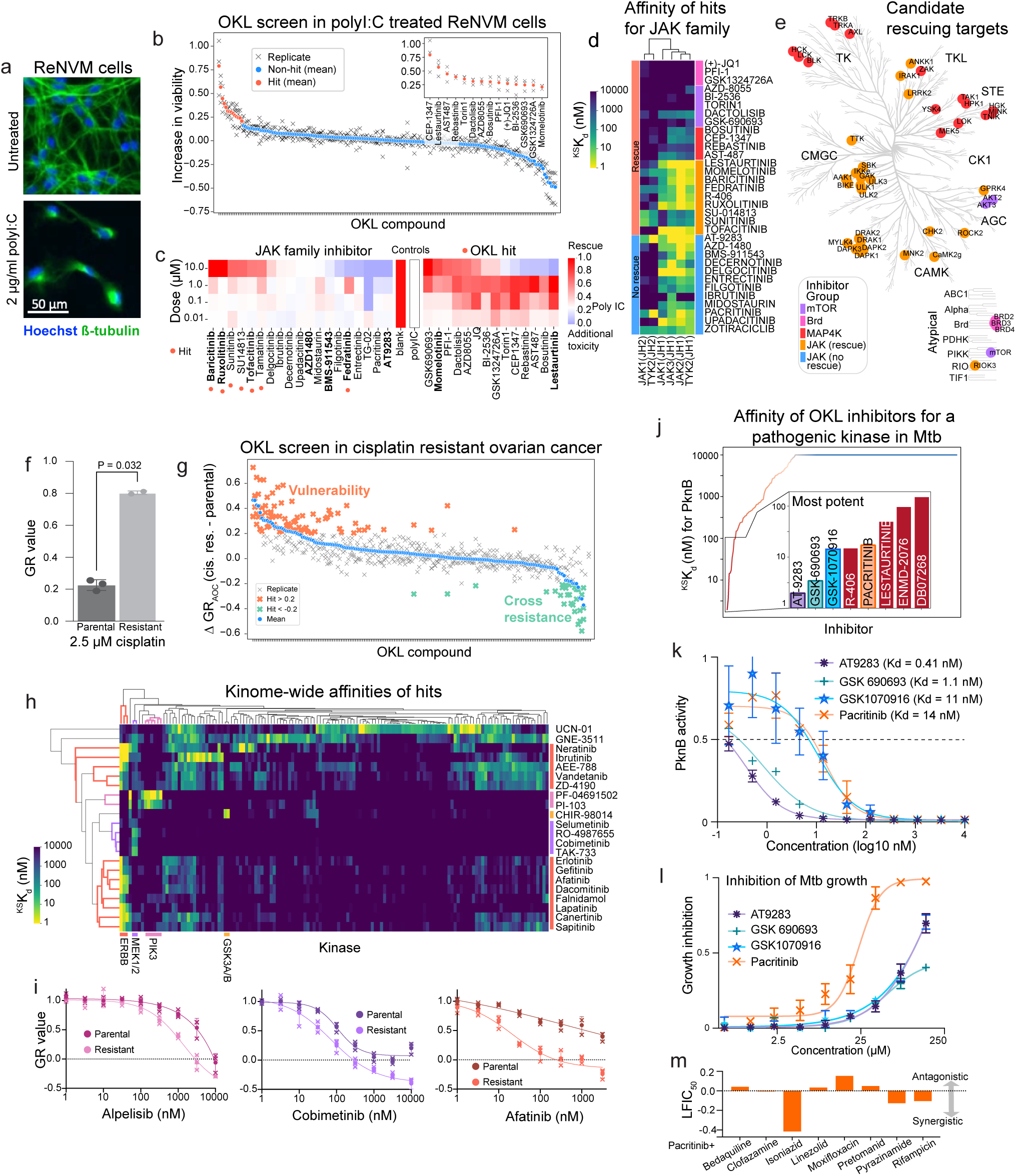
OKL use cases. **(a)** Images of ReNcell® VM cells under control conditions (top) and treated with 2 µg/ml polyI:C (bottom) after seven days. Nuclei in blue and ß-tubulin in green; scale bar 50 µm. **(b)** Change in percent viability conferred by the addition of each OKL compound at 1 µM to polyI:C treated cells (relative to polyI:C alone). The means of biological triplicates (denoted with x’s) are shown as solid o’s in red for hits, and blue for all others. **(c)** Heatmap showing percent viability of ReNcell® VM cells treated with polyI:C and increasing concentrations of JAK inhibitors, and the OKL hits identified in (b). JAK inhibitors that conferred an increase in viability ≥ 20% are denoted with orange dots. **(d)** ^KS^Kd values for the JAK family kinases for all inhibitors shown in (c) arranged by phenotype and common targets. **(e)** Common targets of rescuing agents shown on the kinome tree colored by the groups shown in (d). **(f)** Bar graph showing GR values for parental and cisplatin resistant SNU8 cells treated with 2.5 µM cisplatin for 72 h. **(g)** Difference in GR_AOC_ between cisplatin resistant and parental SNU8 cells for each OKL inhibitor (n=3 biological replicates). Higher GR_AOC_ indicates sensitivity, while lower GR_AOC_ indicates resistance. Individual replicates are shown with x’s, and the mean values with o’s. Hits are denoted by bold orange if more effective and by bold green if less effective in the resistant cells. **(h)** Clustermap of ^KS^K_d_ values for hits that were more effective in resistant cells in at least two primary screens. Only kinases that were inhibited with ^KS^K_d_ ≤ 100 nM by at least one hit are shown. **(i)** Dose response curves for inhibitors of targets identified in (h) for parental and cisplatin-resistant SNU8 cells. **(j)** ^KS^K_d_ values for OKL compounds against PknB, the strongest inhibitors are shown in the inset bar graph. **(k)** 11-point K_d_ curves and K_d_ values for four of the top inhibitors from (j) for PknB. **(l)** Dose-dependent growth inhibition of Mtb treated with the same drugs in the butyrate growth condition. **(m)** Effect of pacritinib in combination with the drugs indicated on Mtb growth.

This yielded 21 rescuing molecules including 9/21 inhibitors whose assigned targets were JAK family kinases. Variability in rescue did not correlate with affinity for assigned JAK targets, which was generally in the nanomolar range (**Figure 5c**). Moreover, other OKL hits did not detectably bind JAK family members (**Figure 5d**). We found that the rescuing agents, but not the non-rescuing agents, had micromolar ^KS^K_d_s for multiple kinases other than JAK1-3/TYK2 including, ULK1-3, the numb associated kinases GAK, AAK1 and BIKE, and the entire DAPK family (DAPK1-3 and DRAK1-2) that function in endocytosis, viral trafficking, and autophagy (**Figure 5e, Supplementary Figure 4a**)^45,46^. This is consistent with recent data showing that numb associated kinases contribute to the activity of baricitinib (Olumiant®), whose assigned targets are JAK1/2, against SARS-COV-2 infectivity, an indication for which baricitinib was recently approved^47^.

The remaining OKL hits fell into three other groups: bromodomain inhibitors (these were in OKL since bromodomains are considered atypical kinase domains) that do not bind any kinases in our analysis; selective AKT/mTOR inhibitors; and promiscuous inhibitors that share affinity for MAP4 kinases (**Figure 5e, Supplementary Figure 4b-c**). Autophagy, the tightly controlled process by which cells selectively degrade and recycle proteins and organelles, is often dysregulated in AD^48^. The finding that combinations of ULK, DAPK, mTOR, and bromodomain inhibition reduces cdsRNA-mediated neuronal death suggests a role for autophagy: autophagy is downregulated by mTOR via ULK1, a key regulator of autophagy initiation^49^ and BRD4 via transcriptional control of autophagy and lysosome related genes^50^. Inhibition of MAP4 kinases, and not JAK inhibition, has recently been shown to promote neuroprotection against paclitaxel-induced peripheral neuropathy^51^. MAP4 kinases are also the assigned targets of prosetin^52^, a drug currently in clinical trials for Amyotrophic Lateral Sclerosis (ALS). Together, these results suggest that MAP4 kinases are a node of convergence for multiple neurodegenerative conditions. This use case also illustrates the value of extending the OKL dataset in a context specific manner.

### Other applications of OKL-affinity data

The second use case is an example of chemical genetics in which small molecules are used to dissect a cellular phenotype, in this case platinum-resistant high grade serous ovarian cancer (HGSOC). Platinum-based therapies are standard of care for HGSOC and commonly elicit tumor regression that is followed by resistance and recurrence^53^. We established a cell culture model of acquired cisplatin resistance by growing SNU8 cells in the presence of increasing concentrations of cisplatin until significant (but partial) resistance was achieved (**Figure 5f**). Screening parental and resistant SNU8 cells with the OKL at four concentrations (10 µM, 1 µM, 100 nM, and 10 nM) yielded 22 hits (**Figure 5g**); rescreening was performed by collecting nine-point dose response curves in the absence of cisplatin. Thirteen hits were inhibitors of EGFR-family members (including 11 assigned EGFR inhibitors), four were assigned to be MEK inhibitors and two were PI3K inhibitors (**Figure 5h-i, Supplementary Figure 5a**). These data demonstrate a requirement for EGFR signal transduction in cisplatin resistance, fulfilling our “chemical genetics” goal. They are also consistent with previous reports in cells and animal models^54,55^. While EGFR inhibition has not as yet proven to be clinically effective in the treatment of platinum-refractory HGSOC^56^, but inhibition of kinases downstream of EGFR such as MEK or PI3K is another possibility as is inhibition of kinases not in the EGFR pathway (as commonly defined) but inhibited by hits in our screen such as GNE-3511 (a DLK inhibitor), CHIR-98014 (a GSK3 inhibitor), and UCN-01 (a broadly active compound).

In the third use case we looked for inhibitors of non-mammalian (pathogen) kinases present in the KINOMEscan panel. We identified eight inhibitors with ^KS^K_d_ ≤ 100 nM for PKNB (**Figure 5j**), a serine threonine kinase essential for growth in *M. tuberculosis* (Mtb)^57–59^. We confirmed the potency of the top hits with 11-point K_d_ assays and found all had K_d_ < 15 nM (**Figure 5k**). When tested in Mtb growth-inhibition assays, pacritinib achieved half-maximal growth inhibition at 22.4 µM (95% confidence interval 19.9 µM – 25.3 µM) when butyrate was the carbon source (**Figure 5l**) but it was not effective under dormancy conditions^60^ (**Supplementary Figure 5b**) consistent with the role of PKNB in Mtb growth. Since Mtb is treated with multiple drugs, we tested pacritinib in combination with eight antibiotics that are core components in standard of care treatment cocktails; we observed substantial synergy with isoniazid, pyrazinamide, and rifampicin (**Figure 5m**). Pacritinib is indicated for treatment of myelofibrosis and thrombocytopenia^61^ and its Cmax value is well below the effective concentration for inhibition of Mtb growth in vitro, but development of a pacritinib-like molecule based on existing medicinal chemistry series may be possible. We also identified 14 potent inhibitors (^KS^K_d_ ≤ 100 nM) for the *P. falciparum* kinase PFCDPK1 and one for PFPK5 (**Supplementary Figure 5c**). PFPK5 is structurally similar to mammalian CDKs^62^ and is most potently bound by PHA703887 whose assigned target is CDK2. This suggests that molecules intended to target human CDKs could be repurposed to develop anti-malaria drugs.

### Recommendations on future KINOMEscans and interpreting existing data

A review of the literature suggests that it is rare to perform KINOMEscan assays at more than one dose, presumably due to cost. Eurofins suggests that hits are those targets that retain <35% of binding relative to a negative control regardless of the kinase identity or compound concentration. To determine whether this is an optimal approach, we compared percent control values at each of our screening concentrations to K_d_s from ChEMBL (**Supplementary Figure 6a**) and evaluated the likelihood of correctly classifying a compound–target interaction across a range of percent control thresholds (see Methods, **Supplementary Figure 6b-c**). We found that the optimal screening concentration and threshold differed based on the objective of the screen and tolerance for false positive as opposed to false negative data (Type 1 and 2 errors; **Figure 6a**). For example, were one to screen ruxolitinib at 10 μM with the goal of identifying targets having K_d_ < 1000 nM, Eurofins standard threshold of 35% would return 94 false positives and two false negatives whereas our recommended cutoff of 22% for this screening scenario returns 64 false positives and two false negatives (**Figure 6b-c**). This reduces false positives (by one-third) but given the amount of effort required to investigate off-targets, we also studied the value of performing KINOMEscans at two concentrations. To do this, we used Bayesian inference to estimate ^KS^K_d_ values when two doses were screened (see **Supplementary Note, Supplementary Figure 6d, Figure 6d**) and found that 10 µM and 100 nM datapoints were optimal and were well correlated with four-point data (R² = 0.97, mean squared error = 0.094 for two dose vs. four dose ^KS^K_d_ values, **Figure 6d-e**). Using multiple doses improves the overall predictive performance for identifying inhibitor-kinase pairs with ^KS^K_d_ < 1 µM as compared to a single dose (the F_1_ score increases from 0.78 to 0.82). We conclude that, in many cases it is likely to be more efficient to collect data on new compounds at two concentrations rather than one, as is the current norm.

**Figure 6:**
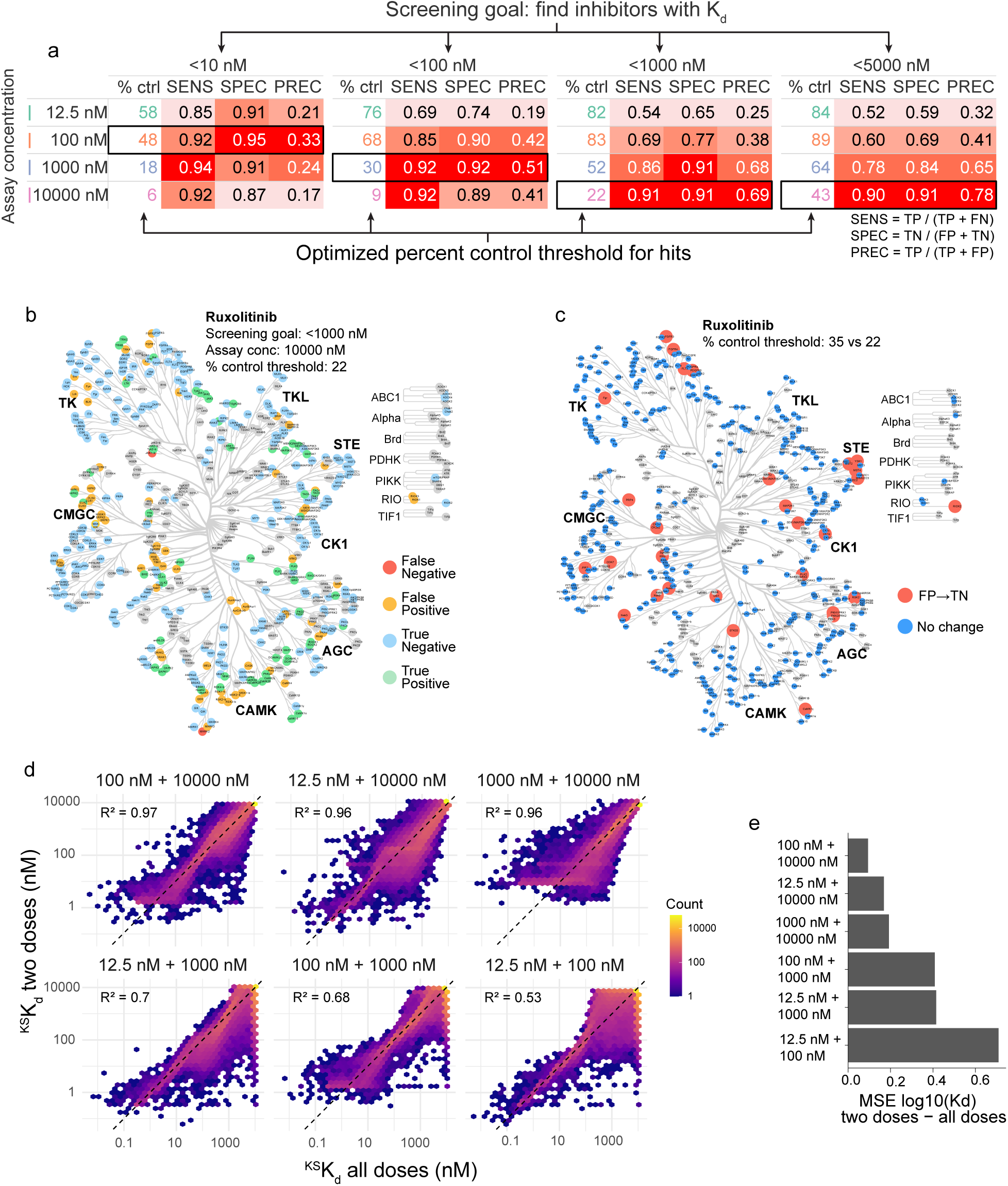
Recommendations for KINOMEscan analysis and interpretation. **(a)** Look-up table based on screening goal and assay concentration showing the sensitivity (SENS), specificity (SPEC) and precision (PREC) for different optimal cutoff values. **(b)** Kinome tree showing the classification of each kinase in a KINOMEscan of 10 µM ruxolitinib using a percent control threshold of 22. **(c)** Kinome tree showing the kinases that are misclassified using a 35% threshold and correctly classified using a 22% threshold. **(d)** Hexagonally binned scatter plots showing the ^KS^K_d_ values estimated from all two dose pairs with respect to those estimated from all four doses. **(e)** Summary barplot of the mean squared error (MSE) of log_10_(K_d_) for all compound–kinase pairs for ^KS^K_d_ values estimated from all two dose pairs.

## DISCUSSION

The biological and clinical activities of molecules used as human therapeutics and tool compounds are commonly described in terms of a limited set of assigned targets, although it is generally understood that a substantial number of additional proteins are also bound. In some cases identifying these other targets can lead to striking changes in therapeutic focus and mechanism of action. Crizotinib, for example, was developed by Pfizer as an inhibitor of the cMET RTK but became a first in class inhibitor of ALK when its additional activities were discovered in cell line studies^63^ and confirmed in clinical trials^64^. It is now possible to profile kinase inhibitors against the great majority of the human kinome using assays such as KINOMEscan, but such data has only sporadically become publicly available, particularly for therapeutics. It has therefore been difficult to draw general conclusions about the magnitude and significance of kinase inhibitor polypharmacology. In this paper we address this by assembling a kinase inhibitor library (OKL) that balances competing demands of compactness, chemical diversity, target coverage, and degree of clinical development. Performing KINOMEscan assays at four doses followed by Bayesian inference of binding constants (^KS^K_d_) yielded data that was well correlated with existing information in ChEMBL while filling in a large number of unknown quantities necessary for systematic analysis.

These data show that many kinase inhibitors, even those used as human therapeutics are potent, but not particularly selective, with a PI_max_ < 0.2, meaning that they bind multiple targets with similar affinities. Strikingly, for kinase inhibitors used as therapeutics, an assigned target is the *highest* affinity binder in only 23% of cases examined. We find no evidence that drugs that have succeeded in clinical development are more selective than those that were abandoned after entering trials. Moreover, the average selectivity of therapeutics has not appreciably increased over time, although the last few years have witnessed the continued development of some highly selective compounds. It does not necessarily follow that kinases inhibited by a particular compound are part of the biological or therapeutic mechanism of action, but it is noteworthy that off-targets include many understudied kinases whose functions are poorly understood. Thus, it is tempting to speculate that it is kinase inhibitor polypharmacology rather than selectivity that explains why this class of drugs has been so successful^65^.

The availability of systematic kinase inhibitor activity data has multiple potential uses and the data in this paper are therefore freely available in raw and processed forms (and have been incorporated in smallmoleculesuite.org). One striking finding is that there exists a subset of kinases that are substantially more promiscuous than the kinome as whole. In some cases, it is possible to rationalize this in terms of known structural properties (e.g. the DFG in and out conformations) but in many others it is not, suggesting that additional structural studies are necessary. Unrecognized patterns of selective binding are also revealing. For example, potent binding of alectinib (assigned target ALK) to the liver-specific phosphorylase kinase isoform PHKG2 may explain the observed hepatoxicity of this compound. Not all potent ALK binders also bind PHKG2, however, suggesting that it might be possible to increase therapeutic index by considering PHKG2 as an anti-target during compound development^66^. Systematic affinity data also reveal multiple cases in which closely related kinases can be differentially targeted with existing chemical backbones, as well as cases in which this is not possible; such information can guide screening and medicinal chemistry efforts. The spectrum of small molecule binding data for each kinase also represents a different way to construct a kinome tree. Currently this is based on sequence alignment and presumed phylogeny, but a tree constructed by clustering small molecule data would be substantially more useful for drug discovery and mechanism of action studies. We estimate that a roughly two-fold expansion of the OKL library to include the latest clinical and pre-clinical traditional kinase inhibitors, PROTACs and molecular glues^67^ would make this possible and is well within the capacity of an academic lab with dedicated funding. The current paper includes computational tools and practical guidelines to guide such an effort, or any other use of KINOMEscan screening, that go beyond what the vendor (Eurofins) recommends.

The possibility of instantiating the OKL in this paper using commercial resources makes it feasible to use it for screening. Moreover, in cases in which an inhibitor with multiple targets induces a desirable phenotype, an OKL screen can help to narrow down the list of functional targets even in settings in which RNAi and similar genetic approaches are challenging. For example, in the work of Petrova et al^51^. a multi-targeting kinase inhibitor, KW-2449, was found to be neuroprotective against paclitaxel-induced peripheral neuropathy and an OKL screen suggested that STE20 kinases play a critical role^51^. In this paper we use OKL to screen for potential neuroprotective agents in a model of Alzheimer’s disease and platinum-resistant ovarian cancer. In each case, the availability of systematic binding data made the screens easier to interpret because they revealed common and, in some cases, unrecognized off-target activities. These studies also demonstrated that it is possible to selectively expand the OKL and associated data to include new molecules focused on specific kinase subsets (in our case JAK family inhibitors).

Drugs targeting kinases are indicated for many conditions, and kinase inhibition remains an area of intense focus in academia and pharma. Despite decades of effort, moving new molecules through development and predicting which are most likely to make good therapeutics remains challenging. The current work challenges some common assumptions about what makes a good human therapeutic, while providing an approach to maximizing the information that can be obtained from phenotypic screens.

## Acknowledgements

We thank ICCB-L for compound management, and Eurofins for helpful discussions during data acquisition. We thank J. Tefft and C. Dessauges for helpful discussions, and manuscript editing, and N. Moret for OKL compound selection. Research reported in this publication was primarily supported by the DARPA PANACEA program, award HR0011-19-2-0022, the NIH Illuminating the Druggable Genome Grant U24-DK116204, the Ludwig Center at Harvard and the Henry and Belina Termeer Foundation. It was also supported by NSF 22-599, EPSCoR RII Track-1, Award Number DQDBM7FGJPC5 (RC), NIH R01AI150684 (BA), and NIH R01 2R01AG058063-06 (MWA). The content is solely the responsibility of the authors and does not necessarily represent the official views of the National Institutes of Health.

## Data availability

All data are included as Supplementary Tables.

## Author Contributions

Conceptualization: CEM, CH and PKS; Methodology: CEM, CH, BMG, KAS, NC, CV, MC, HD, MPM, SR, and RC; Investigation: CEM, CH, CV and MC; Formal analysis: CEM, CH and BMG; Resource: CEM, CH and PKS; Visualization: CEM, CH and RC; Software: CH, SR, BMG; Funding acquisition: BA, RC and PKS; Supervision: CEM, BA, MWA, RC, BMG and PKS; Writing – original draft: CEM, CH and PKS; Writing – review & editing: CEM, CH, BMG and PKS.

## METHODS

### KINOMEscan and enzymatic kinase activity assays

Small molecule kinase inhibitors were purchased from commercial vendors (**Supplementary Table 1**). Stock solutions were prepared in dDMSO at 10 mM for all compounds with the following exceptions due to limitations in solubility: CAY10561 at 5 mM, CHIR-98104 at 2 mM, gedatolisib at 4 mM, and indirubin at 3 mM. Inhibitors were arrayed in 96 well plates, 100 µl per inhibitor, and sent to Eurofins for KINOMEscan scanMAX profiling. scanMAX profiling was performed in two batches: in the first batch, scans were performed at 10 µM and 100 nM; scans in the second batch were performed at 1 µM and 12.5 nM concentrations. A subset of inhibitors was only screened in the first batch and we therefore only have data points at 10 µM and 100 nM. For each concentration of each inhibitor, a percent of control value was returned for 406 WT kinases and 59 mutant kinases and three non-mammalian kinases (**Supplementary Table 2**) for analysis.

For enzymatic assays used for confirmatory studies, kinase inhibitors were sent to Reaction Biology for follow-up IC50 assays, performed in duplicate at 10-dose points (HotSpot™ radiometric assays, with 10 µM ATP), or to Eurofins for follow-up K_d_ assays performed in duplicate at 11-dose points (KdELECT®) depending on assay availability.

### Inhibitor metadata

Inhibitor type assignments were sourced from the literature as much as possible. For inhibitors with K_d_ < 3 µM for any ABL1 target, the relative affinity of kinase inhibitors for WT or mutant ABL1 in its nonphosphorylated vs. phosphorylated states which has been previously shown to distinguish type I (no preference for activation state) from type II inhibitors (preference for the nonphosphorylated state)^7,68^ was calculated. Assigned targets for each inhibitor were sourced from the literature and from the vendors.

### Quality control of KINOMEscan data

Quality of the dataset was assessed by plotting the percent control values in four-point dose response curves for every inhibitor-kinase pair and classifying their shapes. Eurofins recommends using a cutoff of 35% control for hits. Based on this we defined five classes: *non-binding*, no measurements below 35% control; *binding*, two or more measurements below 35% control, all measurements concordant; *weakly binding (high confidence)*, one measurement below 35%, p<0.1 for a linear model fit to the dose response curve; *weakly binding (low confidence)*, none of the above apply (these are usually curves where the 10 µM data point is the only data point below 35%); and *discordant*, one measurement below 35% control, one measurement, at a higher concentration, above 35% and at least two-fold higher than a previous value (**Supplementary Figure 1a**). We found that 73% of the dose responses measured were classified as non-binding. 2% of the ∼90k dose response relationships were classified as discordant, accounting for 6% of the data points when non-binding curves are excluded (**Supplementary Figure 1b**). A subset of inhibitors and kinases accounted for a disproportionate number of discordant curves suggesting that there were systematic errors in the dataset (**Supplementary Figure 1c, d**). The dose responses for these kinases and inhibitors showed clear batch effects (the percent control values for the 1 µM and 12.5 nM data points were often lower than the 10 µM and 100 nM data points) (**Supplementary Figure 1e, f**, left panels). In scanMAX assays, multiple kinases are tested in a single well so a technical error (such as a liquid handling mistake) would affect the data for several kinases – it is likely that the kinases with the most discordant dose response curves were multiplexed during data collection. Therefore, the assays for all inhibitors for the seven worst performing kinases (**Supplementary Figure 1d**), and for the 12.5 nM and 1000 nM concentrations of erlotinib (the worst performing inhibitor) were repeated. The concordance of the dose responses for the conditions that were repeated improved substantially (**Supplementary Figure 1e, f**, right panels) and the overall fraction of discordant dose responses was reduced to 1% of the total dataset (**Supplementary Figure 1g**). The remaining discordant dose responses were randomly distributed across kinases and inhibitors and were removed from all analyses. There were an additional 300 data points that were initially reported as 100% control (no binding) that were qPCR dropouts (Eurofins provided this list, but it is not routinely shared). These data points were also excluded.

### KINOMEscan-derived K_d_ values (^KS^K_d_), target affinity spectra

To estimate dissociation constants (K_d_) and Hill slopes from KINOMEscan dose-response data, we implemented a Bayesian hierarchical model that is described in detail in the **Supplementary Note**. To facilitate comparisons with previous analyses, we converted the ^KS^K_d_ values to target affinity spectra (TAS): ^KS^K_d_ > 10 µM corresponds to a TAS value of 10; ^KS^K_d_ between 1 µM and 10 µM to TAS 3; ^KS^K_d_ between 100 nM and 1 µM to TAS 2; and ^KS^K_d_ < 12.5 corresponds to TAS 1.

### Selectivity metrics

We calculated the partition index^21,69^ (PI) for a given inhibitor with target 𝑖 using our ^KS^K_d_ values (equation 3).

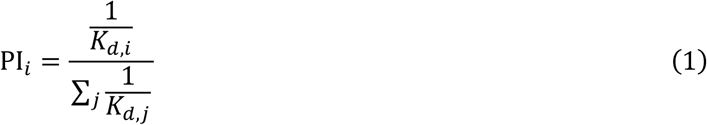

To identify potential chemical probes, we used the Concentration and Target Dependent Selectivity (CATDS) metric^6^, which has previously been applied for this purpose^10^. Specifically, we computed CATDS_most-potent_, which quantifies the proportion of a compound’s total target engagement attributed to its most potent target, evaluated at the concentration equal to the K_d_ of that target. For each compound, we identified the most potently bound target as the kinase with the lowest fitted ^KS^K_d_ value (𝐾_d,ref_). We then calculated the predicted target engagement 𝐸*_i_* for every measured target 𝑖 at a compound concentration equal to 𝐾*_d_*_,ref_, using the fitted 𝐾*_d,i_* and Hill slope ℎ*_i_*:

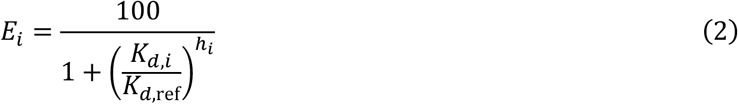

CATDS_most-potent_ for the highest affinity target is then the fraction of total predicted engagement accounted for by that target:

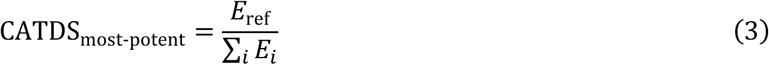

### ChEMBL binding data

Data were obtained from the ChEMBL PostgreSQL database (version 33) and filtered based on standard unit types (K_d_, K_d,app_, and K_i_) and assays measuring direct protein targets, excluding cell- and tissue-based assays. Agreement between ^KS^Kd estimates and half-maximal inhibitory concentrations (IC_50_) in ChEMBL was R = 0.78 reflecting differences in what each metric represents; comparisons were therefore limited to Kd data in ChEMBL. However, binding constants derived from multiplexed inhibitor bead assays only correlated weakly (R = 0.66) with other Kd data in ChEMBL and were therefore also excluded when we compared our ^KS^K_d_ results and single dose data to ChEMBL K_d_s (**Figure 1f, Supplementary Figure 6a**).

### OKL screening in ReNcell® VM cells

ReN VM cells were maintained free of mycoplasma in their recommended growth conditions in ReNcell® NSC maintenance media (EMD Millipore, Danvers MA), supplemented with 20 ng/ml rhEGF (Millipore Sigma, Burlington, MA), 20 ng/ml FGF (Millipore Sigma, Burlington, MA); 5% CO_2_, 37°C, on growth factor reduced Matrigel (Corning, Corning, NY). Prior to drug treatments, cells were differentiated on 10 cm plates by removing growth factors from the media for seven days. Cells were then removed from the plates using accutase (EMD Millipore, Danvers, MA), resuspended, plated in 96 well plates and allowed to differentiate for another seven days. Media was changed twice per week during differentiation. For OKL screening (final concentration 1 µM), drug treatments were added by acoustic transfer using an ECHO 655 (Beckman Coulter, Brea, CA) at the ICCB-L screening facility one hour prior to the addition of 2 µg/ml (final concentration) of poly(I:C) HMW (Invivogen, San Diego, CA). After four (replicate 1) or six (replicates 2 and 3) days, an equal volume of Cell Titer Glo 2.0 (Promega, Madison, WI) was added to each well, incubated at room temperature with rocking for 10 min. and luminescence was measured with a Synergy H1 (BioTek, Winooski, VT) plate reader. Validation experiments were performed in the same way except drug treatments were delivered with a D300 (Tecan, Morrisville, NC) digital drug dispenser, and data were collected after seven days.

### Dose response measurements in ovarian cancer cells

SNU8 ovarian cancer cells were maintained in RPMI-1640 supplemented with 2 mM L-glutamine, 25 mM HEPES, 10% fetal bovine serum and 1% penicillin/streptomycin at 37°C and 5% CO2. Cells were identity validated by short tandem repeat profiling^70^ and verified to be free from mycoplasma. Cisplatin-resistant cells were generated by adding cisplatin to the growth media starting at 0.5 µM. The dose was escalated incrementally, as the cells allowed, until they were stable in the presence of 3 µM cisplatin (approximately six months). For all dose response measurements, the parental and resistant cells were plated in 384 well CellCarrier ULTRA plates (Perkin Elmer, Waltham, MA) at 750 cells per well in 60 µl media, in the absence of cisplatin. Cells were allowed to adhere for 24 h prior to drug treatment. Drug treatments (final concentrations of 10, 1, 0.1, and 0.01 µM) were delivered via pin transfer with a custom E2C2515-UL Scara robot (Epson, Long Beach, CA) coupled to stainless steel pins (V&P Scientific, San Diego, CA) from 1000x library plates at the ICCB-L screening facility at Harvard Medical School for the screens, and using a D300 digital drug dispenser in nine-point half-log dilution series for validation experiments (Tecan, Morrisville, NC). At the time drugs were added, a time = 0 plate was stained and fixed using the Deep Dye Drop protocol^71^. Following six days in drug, all remaining assay plates were stained and fixed using the Deep Dye Drop protocol. In brief, live cells were stained with LIVE/DEAD Red (LDR) (Thermo Fisher, Waltham, MA), and pulsed with EdU (Lumiprobe, Waltham, MA) for one hour in a solution of 10% Optiprep™ (a density gradient medium) (Sigma, St. Louis, MO) in phosphate buffered saline (PBS). Cells were then fixed with 4% formaldehyde prepared in 20% Optiprep™ for 30 min. The addition of sequentially denser solutions enables staining and fixation without the need for any aspirate or wash steps thus preserving cells susceptible to loss, reducing the number of protocol steps, and sparing reagents and costs^71^. Following fixation, cells were permeabilized with 0.5% Triton X-100 for 15 min., the EdU was labeled with cy3-azide (Lumiprobe, Waltham, MA) using Click chemistry for 30 min., cells blocked in Odyssey buffer (Li-COR Biosciences, Lincoln, NE) for one hour at room temperature, and finally stained overnight with Hoechst 33342 (1:5000, Thermo Fisher, Waltham, MA) and an Alexa 488 conjugated anti-phospho histone H3 antibody prepared in Odyssey (1:2000, Cell Signaling Technologies, Danvers, MA).

Stained and fixed cells were imaged with an ImageXpress Micro-Confocal (IXM-C) high throughput microscope equipped with a plate hotel and robotic arm (Molecular Devices, San Jose, CA). Four fields of view were imaged per well with a 10x objective for full well coverage. Image analysis was performed with MetaXpress software (version 6.5.3.427, Molecular Devices, San Jose, CA): nuclei were segmented, and nuclear masks were defined based on the Hoechst signal. The nuclear mask was dilated to create a ring mask, and the intensity of each signal in each mask was measured. Local background was accounted-for by subtracting the intensity of each marker in the ring mask from the nuclear mask. Objects touching the edge of the images were excluded from analysis. Analysis of the raw feature data was performed using custom scripts (https://labsyspharm.github.io/dye-drop-microsite/)^71^. The integrated Hoechst intensity was used for DNA content quantification, EdU intensity for S-phase cells, and phospho-histone H3 intensity for M-phase cells. LDR signal was used to identify dead cells. Growth rate inhibition (GR) values and metrics were calculated as defined by the number of viable cells at the time of drug treatment (t=0), and at endpoint under untreated and treated conditions^72^. A compound was considered to induce a stronger effect in the cisplatin resistant cells if the difference between the area over the GR curve (GR_AOC_) was at least 0.2 lower than for the same treatment in the parental controls. Compounds that met this condition in at least two biological replicates of the screen were considered hits.

### Molecular Dynamics Simulations

Molecular dynamics (MD) simulations were performed to investigate the structural and energetic basis of CLK-inhibitor interactions for both apo and holo conformations of CLK1 (PDB 6Q8K), CLK2 (PDB 6FYI), CLK3 (PDB 6RCT), and CLK4 (PDB 6FYV) with eight inhibitors (64 simulations in total) using the GROMACS^73,74^ molecular simulation platform. We constructed two system configurations: (1) *apo-CLK systems*, where the inhibitor was positioned near CLK but outside the binding pocket, and (2) *holo-CLK systems*, where the inhibitor was positioned within the CLK binding pocket. We applied the charmm36 force field^75^ to model the systems and parameterized the inhibitors using the Automated Topology Builder (ATB)^76^. To prepare the initial system configurations, we solvated CLK-inhibitor in a cubic simulation box with the TIP3P water model and added Na^+^ or Cl^-^ ions to neutralize the charge and avoid artifacts in long-range electrostatics calculations. We carried out energy minimization using the steepest descent method^77^, followed by a 100-ps equilibration phase under the canonical ensemble (NVT) at 300 K, where we restrained the protein’s heavy atoms. We then carried out isothermal-isobaric equilibration (NPT) at 300 K and 1 bar using the Parrinello-Rahman barostat^78^. Finally, we conducted production MD simulations for 500 ns under the same thermodynamic conditions, using a time step of 2 fs. All equilibration and production runs used the leapfrog integrator^79^. We imposed periodic boundary conditions in all dimensions and calculated long-range electrostatics using the Particle Mesh Ewald (PME) method^80^. To ensure consistency, the same force field parameters and simulation protocols were applied across all systems. Predicted scores were normalized to the highest absolute value of all predictions.

### Dose response measurements in *M. tuberculosis*

Aliquots of pacritinib, AT9283, GSK690693, and GSK1070916 (200 µl at 10 mM in DMSO) were sent to the Aldridge lab at Tufts University. Dose responses for pacritinib, AT9283, GSK690693, and GSK1070916 were measured in *M. tuberculosis* (Mtb, Erdman strain) in butyrate and dormancy growth conditions as previously described^60^.

## Supplementary Note

### Bayesian Estimation of Dissociation Constants from two and four-dose KINOMEscan Data

The Eurofins KINOMEscan platform provides a cost-effective competitive binding assay for profiling compound affinity across ∼75% of the human kinome. We presume that kinases not in the panel cannot be expressed in recombinant form in the T7 phage display or HEK 293 cell system used by Eurofins, although no specific information is publicly available. Conventionally, KINOMEscan data are assessed at a single concentration using a threshold of <35% control remaining to classify a compound–kinase pair as a ‘hit.’ While straightforward, this binary classification discards quantitative information about binding affinity and provides no estimate of the dissociation constant (K_d_). We therefore profiled each of the 192 OKL compounds at four concentrations (12.5, 100, 1000, and 10,000 nM), generating 89,856 four-point dose-response curves. To optimize the extraction of K_d_ vlues from such dose-response data, we utilized a Bayesian inference framework that fits a sigmoidal dose-response model to the observed percent control values, yielding posterior distributions over K_d_ and Hill slope for each compound–kinase pair.

### Model Specification

We model the relationship between inhibitor concentration and the observed percent of control remaining using a four-parameter Hill equation. Let 𝑐 denote the compound concentration, 𝐾_*d*_ the dissociation constant, and ℎ the Hill slope coefficient. The mean predicted response is:

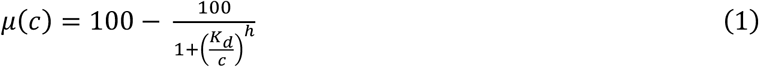

This formulation represents the percent of kinase activity remaining as a function of inhibitor concentration: at low concentrations relative to K_d_, the predicted response approaches 100% (no inhibition), and at high concentrations it approaches 0% (complete inhibition). Measurement error was modeled as heteroscedastic (meaning that the variance in model residuals is not constant across all estimated K_d_, values); specifically, we found that the standard deviation was proportional to the predicted response. We employed an empirically derived error model based on technical reports from Eurofins:

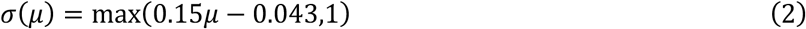

This captures the observation that measurement variability increases with higher percent control values while maintaining a minimum error floor of 1% control. The observed percent control values 𝑦*_i_* at each dose 𝑑*_i_* were modeled as normally distributed around the predicted mean with standard deviation given by the error model.

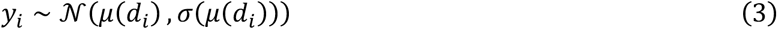

### Prior Distributions

We specified weakly informative priors to constrain parameters to biologically plausible ranges while allowing the data to dominate inference. For the dissociation constant, we placed a log-normal prior centered at 1 μM with wide variance:

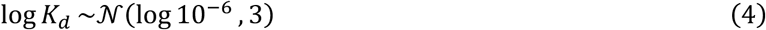

This prior spans a broad range from sub-nanomolar to millimolar affinities, accommodating the full spectrum of biologically relevant dissociation constants. For the Hill slope, we used:

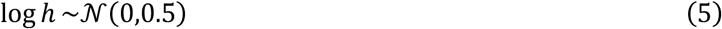

corresponding to a log-normal prior centered at a Hill slope of 1, consistent with standard single-site binding kinetics, while allowing moderate deviations.

### Posterior Inference

For each compound–kinase pair, we sampled from the posterior distribution using the No-U-Turn Sampler (NUTS) as implemented in the nutpie package (https://github.com/pymc-devs/nutpie) with PyMC (version 5.25.1)^69^. We ran four independent Markov chains, each with 1,000 tuning iterations and 2,000 sampling iterations, yielding 8,000 posterior samples per pair. Convergence was assessed using the Gelman–Rubin statistic (R^), with R^ < 1.01 considered indicative of adequate convergence. Point estimates for K_d_ and Hill slope were obtained as posterior medians, and uncertainty was quantified using posterior standard deviations and 95% highest density intervals (HDIs). The full implementation is available as a Python command-line tool (fit_kd_cli.py; github.com/labsyspharm/okl-analysis) that supports parallel execution across compound–kinase pairs with results stored in a SQLite database for reproducibility and seamless continuation after interrupted runs.

### Validation Against ChEMBL Reference Data

To assess the accuracy of our Bayesian K_d_ estimates (denoted ^KS^K_d_), we compared them to K_d_ values deposited in ChEMBL for overlapping compound–kinase pairs where full dose-response curves had been collected using orthogonal assay platforms. We observed good agreement between the four-dose KINOMEscan-derived K_d_ values and the ChEMBL reference values (R² = 0.91; Figure 1f), demonstrating that four concentration points spanning four orders of magnitude are sufficient to estimate dissociation constants with high fidelity using the Bayesian framework.

### Minimum Number of Doses Required for Reliable K_d_ Estimation

Given the cost of kinome-wide profiling, we investigated whether fewer than four concentration points could yield reliable K_d_ estimates. We systematically evaluated all six pairwise combinations of the four assay concentrations (12.5, 100, 1000, and 10,000 nM) by fitting the Bayesian model to each two-dose subset and comparing the resulting K_d_ estimates to those obtained from the full four-dose data (**Figure 6d, e**). As an illustrative example, **Supplementary Figure 6d** shows dose-response fits for lenvatinib binding to EPHB6 across all dose combinations. The four-dose fit (all doses) yields a K_d_ of 35 nM with a narrow 95% HDI. Two-dose subsets that include concentrations spanning the widest dynamic range—particularly 100 nM + 10,000 nM—produce K_d_ estimates (41 nM) and credible intervals closely matching the four-dose fit. In contrast, subsets comprising closely spaced concentrations (e.g., 12.5 nM + 100 nM) yield wider credible intervals and slightly different point estimates, reflecting the reduced information content of narrowly spaced doses.

To quantify the accuracy of K_d_ estimation across the entire dataset, we compared the two-dose and four-dose K_d_ estimates using hexagonally binned scatter plots and computed the R², showing that certain assay concentration combinations perform better for different ranges of K_d_ estimation (**Figure 6d**). To quantify the estimation error when going from four to two doses, we computed mean squared error (MSE) of log_10_(K_d_) for all compound–kinase pairs (**Figure 6e**). The two dose combination of 100 nM and 10,000 nM achieved the best overall performance (relative to a four-dose ground truth), with R² = 0.97 and MSE = 0.094. This was closely followed by 12.5 nM + 10,000 nM (R² = 0.96, MSE = 0.168) and 1000 nM + 10,000 nM (R² = 0.96, MSE = 0.193) (**Figure 6d-e**, **Supplementary Table 9**). Dose pairs that did not include the highest concentration (10,000 nM) performed substantially worse, with MSE values 2–7-fold higher.

The strong performance of the 100 nM + 10,000 nM combination can be understood intuitively: these two concentrations bracket the K_d_ values of most compound–kinase pairs in the dataset, providing information about both the upper and lower plateaus of the dose-response curve.

Concentrations that are both above or both below the K_d_ provide redundant information about one plateau, yielding poorly constrained estimates. Based on these results, we recommend that future KINOMEscan experiments adopt a minimum two-dose design at 100 nM and 10,000 nM when full four-dose profiling is not feasible due to cost constraints.

### Advantages of the Bayesian Approach

The Bayesian framework described here offers several advantages over conventional threshold-based analysis of KINOMEscan data. Most fundamentally, each K_d_ estimate is accompanied by a full posterior distribution, from which 95% highest density credible intervals are derived. This enables downstream analyses to propagate uncertainty rather than relying on point estimates alone, providing a principled measure of confidence for each compound–kinase interaction. The weakly informative priors further serve as a form of regularization, shrinking K_d_ estimates derived from noisy or sparse data toward biologically plausible values. This is particularly important for compound–kinase pairs near the limit of detection, where conventional curve-fitting approaches may yield unstable or biologically implausible estimates.

Because the model encodes the expected shape of the dose-response relationship through the Hill equation and prior distributions, it can also estimate K_d_ values that fall outside the range of tested concentrations. Compounds with K_d_ < 12.5 nM (below the lowest tested concentration) or K_d_ > 10,000 nM (above the highest) can still be assigned meaningful, albeit less certain, affinity estimates, with broader credible intervals that appropriately reflect the increased uncertainty. This capacity for extrapolation has practical consequences for selectivity analysis: quantitative K_d_ estimates across the full affinity range enable the computation of continuous selectivity metrics such as the partition index (PI), which represents the fraction of compound bound to each kinase in a theoretical equimolar mixture. By leveraging the full range of estimated affinities, including sub-nanomolar K_d_ values extrapolated below the lowest assay concentration, the partition index substantially improves the resolution of selectivity profiles relative to threshold-based hit calling, which treats all sub-threshold interactions as equivalent.

For comparison to an alternative non-Bayesian approach, we calculated K_d_ values for each concordant dose response that crossed 50% control by interpolating from a linear relationship between the data points on either side of the 50% mark. We assigned curves that never reached 50% control K_d_ > 10 µM (highest screening dose), and those that were below 50% control at all screening concentrations K_d_ < 100 nM or K_d_ < 12.5 nM depending on the lowest dose screened for each inhibitor. While the correlation between these K_d_ estimates and data in ChEMBL was good (R=0.88), we could not estimate relative affinities below the lowest screening concentration (**Supplementary Figure 1h**). Answers to common questions such as (i) what is the highest affinity target of an inhibitor? and (ii) what target is most selectively bound by an inhibitor? were difficult to answer under these circumstances since they were multi-way ties. For example, a potent multi-targeting inhibitor like dasatinib, inhibits 48 kinases with K_d_ < 12.5 nM, resulting in a 48-way tie for PI_max_. With the Bayesian approach described here, we can see that dasatinib most selectively inhibits EPHA2 with PI_max_ = 0.32 and ^KS^K_d_ = 0.035 nM.

Finally, as demonstrated above, the Bayesian framework yields reliable K_d_ estimates from as few as two concentration points, enabling significant cost reductions without substantially compromising data quality. This makes kinome-wide profiling more accessible and scalable for large compound collections. Notably, numerous past KINOMEscan experiments have already been performed at two doses; these existing datasets could benefit from reanalysis using the approach described here.

## Supplementary Information

**Supplementary Figure 1:**
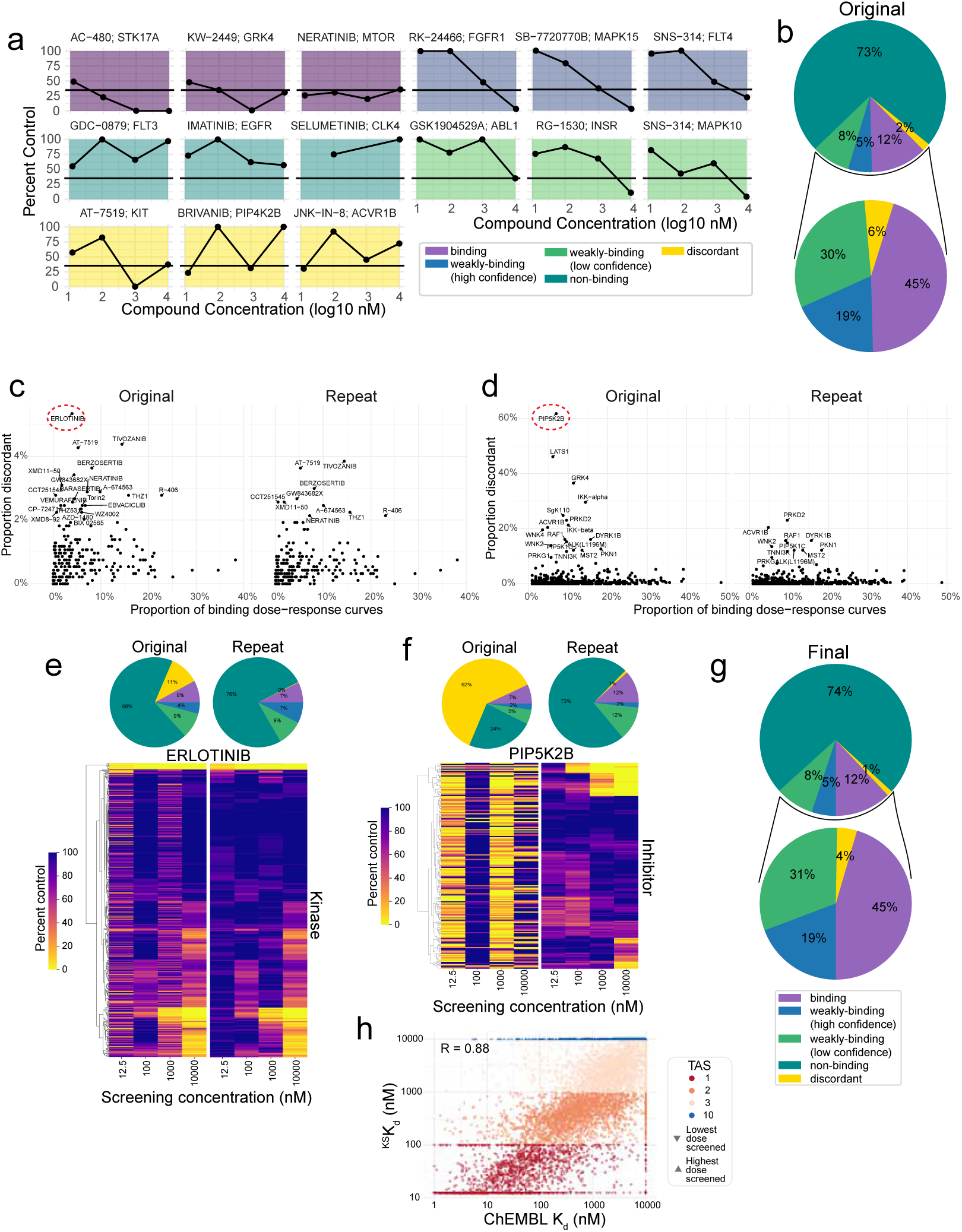
**(a)** Examples of percent control dose response curves for each class as indicated. Discordant dose responses were excluded from the dataset. **(b)** Pie charts showing the fraction of dose responses that fall into each class, and the distribution of dose response classes when the confirmed non-binding class is excluded. **(c)** Scatter plots showing the fraction of discordant dose response curves with respect to the fraction of binding dose response curves for all inhibitors and **(d)** for all kinases in the original dataset (left plots) and in the final post-QC dataset (right plots). **(e)** Heatmaps showing % control values for erlotinib against all kinases at all concentrations tested, and pie charts summarizing the fraction of dose responses in each class pre- and post-QC. **(f)** Heatmaps showing % control values for PIP5K2B and **(g)** Pie charts showing the fraction of dose responses that fall into each class, and the distribution of dose response classes when the confirmed non-binding class is excluded. **(h)** Scatterplot comparing ^KS^K_d_ values estimated by linear interpolation to those previously available in ChEMBL. Data points are colored by TAS value, and the values for each compound’s assigned target(s) are outlined in black.

**Supplementary Figure 2:**
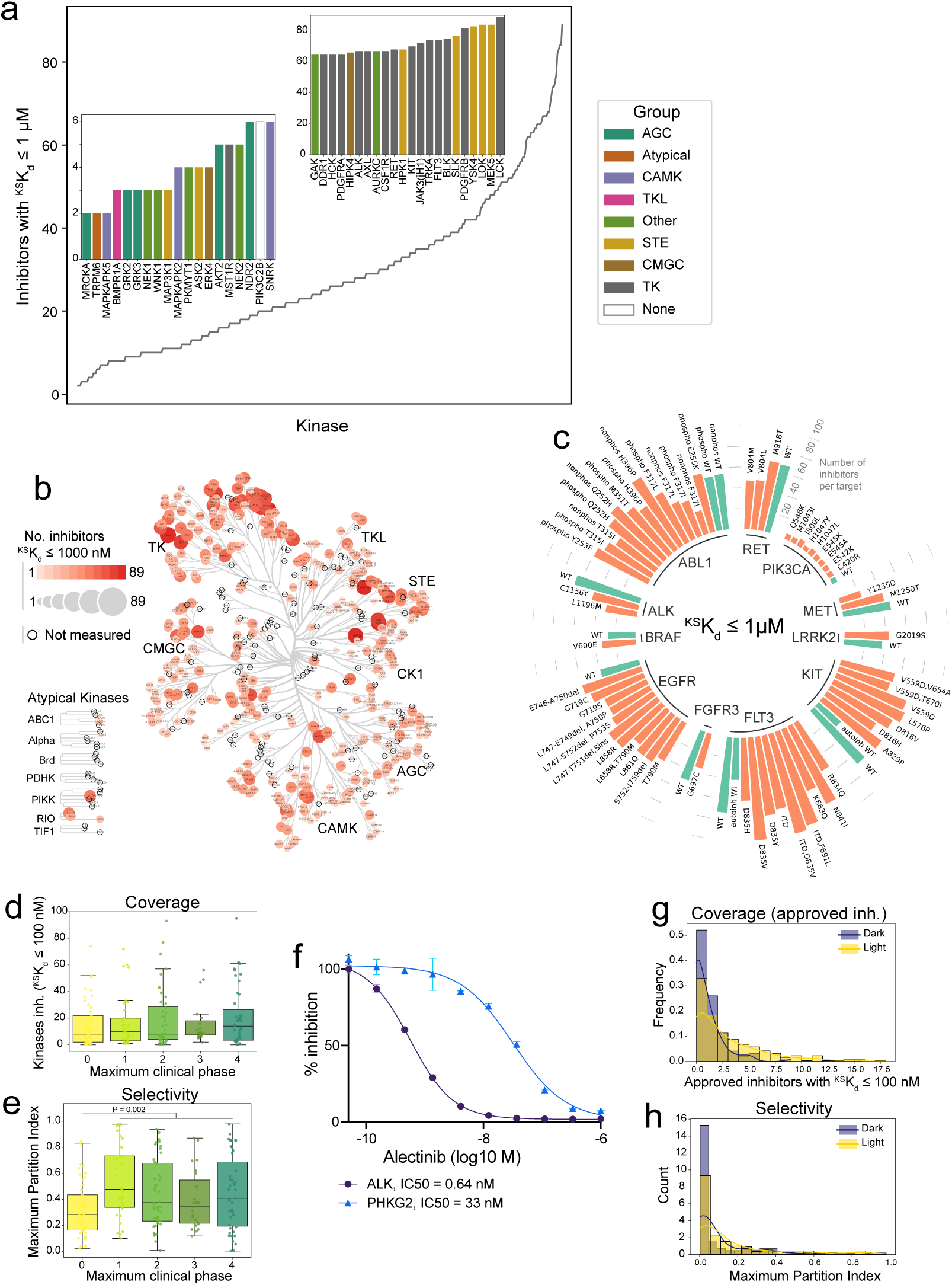
**(a)** The number of inhibitors with ^KS^K_d_ ≤ 1 µM per kinase. The inset plots show the kinases most (right) and least (left) inhibited by OKL compounds colored by kinase group. **(b)** The number of OKL inhibitors (^KS^K_d_ < 1 µM) per kinase shown on the kinome tree by the size and intensity of the markers. **(c)** Radial bar plot showing the number of OKL inhibitors with ^KS^K_d_ ≤ 1 µM for all mutant kinases (red bars) assayed in the KINOMEscan panel. WT kinases are indicated with blue bars. **(d)** Boxplot showing the number of kinases per inhibitor with ^KS^K_d_ ≤ 100 nM by maximum stage of clinical development reached. **(e)** Boxplot showing the maximum partition index for each OKL inhibitor by maximum stage of clinical development reached. P-value is from a one-way ANOVA with Tukey’s correction for multiple comparisons. **(f)** Dose response curves and IC_50_ values for for inhibition of ALK and PHKG2 by alectinib. **(g)** Histogram of the number of approved OKL inhibitors that bind each dark and illuminated kinase with ^KS^K_d_ ≤ 100 nM. **(h)** Histogram of the maximum partition index for dark and illuminate kinases. KDE lines are overlaid for visualization only.

**Supplementary Figure 3:**
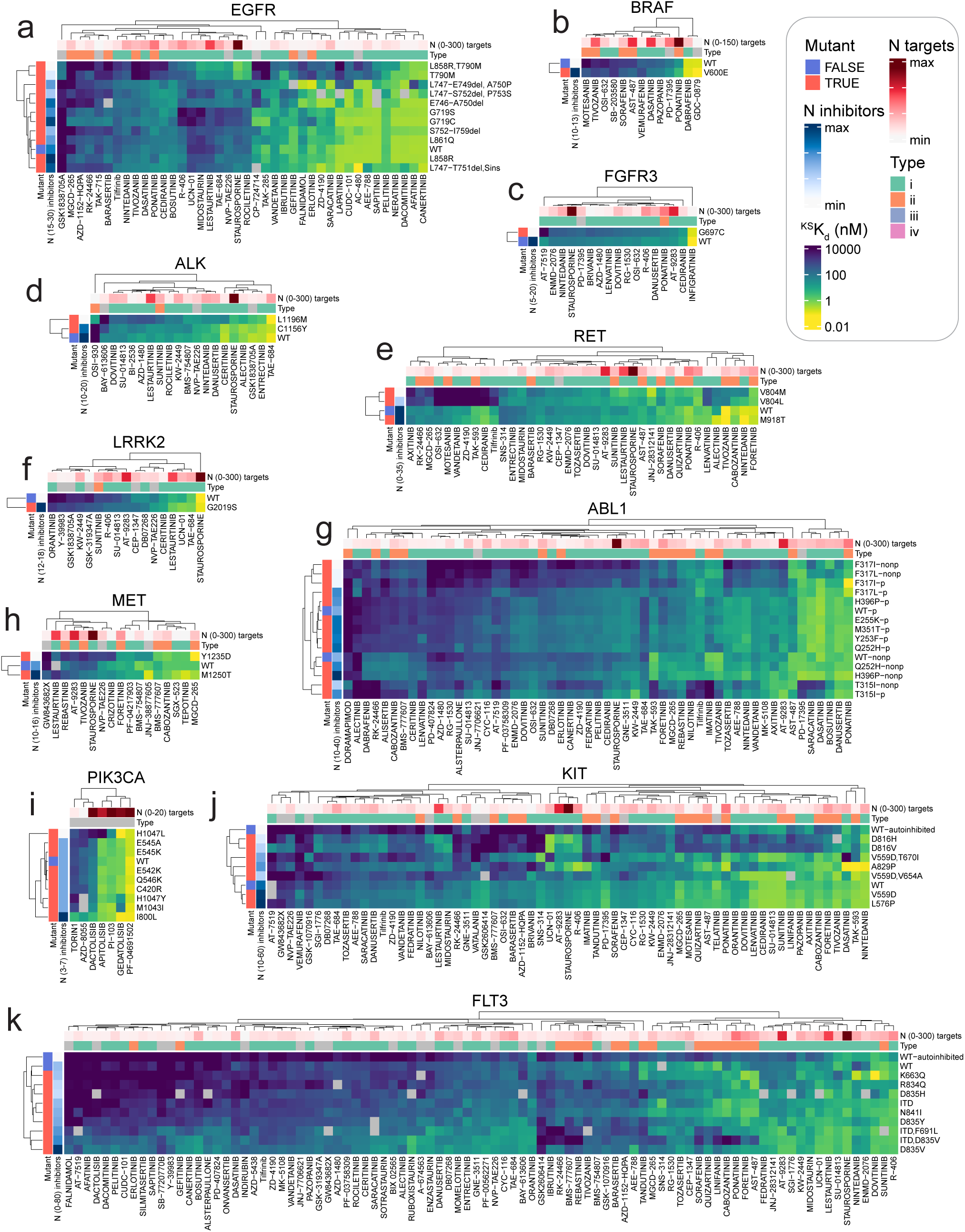
**(a)** Heatmaps with hierarchical clustering showing ^KS^K_d_ values for OKL inhibitors against all variants of EGFR, **(b)** BRAF, **(c)** FGFR3, **(d)** ALK, **(e)** RET, **(f)** LRRK2, **(g)** ABL, **(h)** MET, **(i)** PIK3CA, **(j)** KIT, and **(k)** FLT3 that were assayed.

**Supplementary Figure 4:**
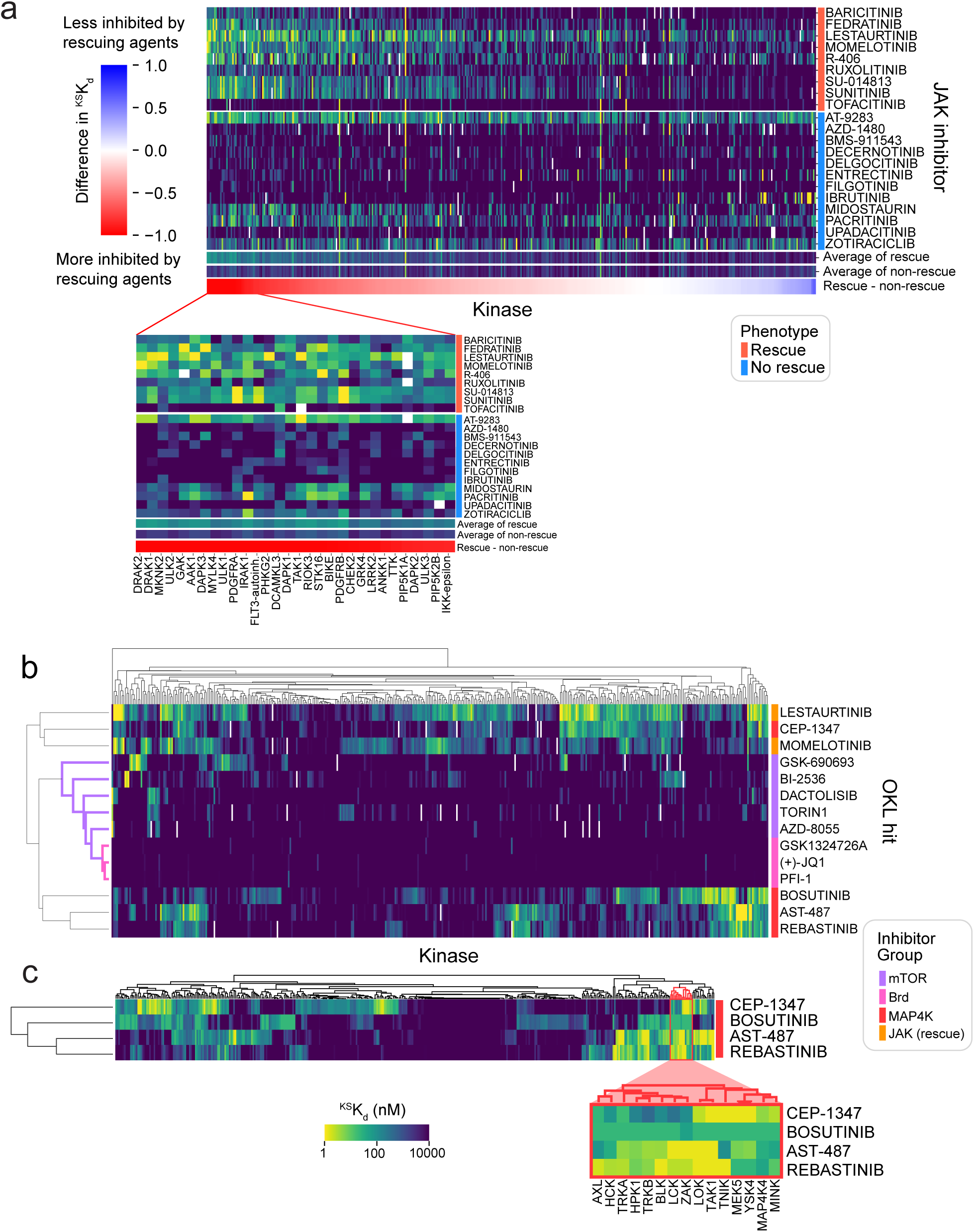
**(a)** Heatmap of ^KS^K_d_ values for nominal JAK family inhibitors. Rows are sorted by phenotype; columns are sorted by the difference in ^KS^K_d_ values between the rescuing and non-rescuing agents. A magnified view of the kinases that are preferentially inhibited by the rescuing agents is provided. **(b)** Clustermap of ^KS^Kd values for OKL compounds that protect ReN VM cells from polyIC. The colored bars represent groupings by targets inhibited. **(c)** Clustermap of ^KS^K_d_ values for the subset of OKL inhibitors denoted with a red bar in (b), and a magnified view of the targets most potently inhibited by those hits.

**Supplementary Figure 5:**
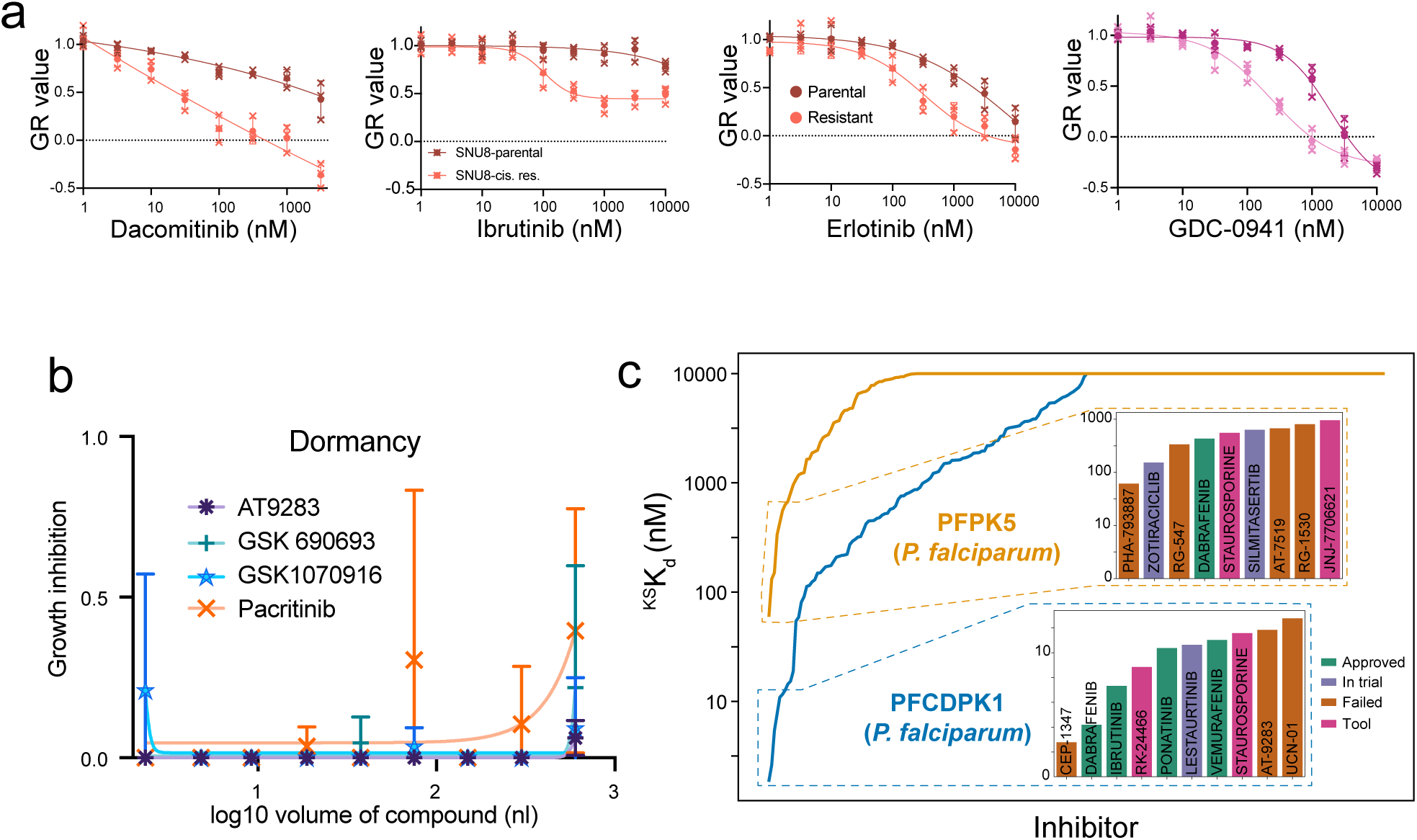
**(a)** Dose response curves for the inhibitors shown in parental and cisplatin-resistant SNU8 cells after six-day treatments. **(b)** Dose-dependent growth inhibition of Mtb treated with the indicated drugs in the dormancy growth condition. **(c)** ^KS^K_d_ values for OKL compounds against the *P. falciparum* (malaria) kinases, PFPK5 (orange) and PFCDPK1 (blue), the strongest inhibitors of each are shown in the inset bar graphs.

**Supplementary Figure 6:**
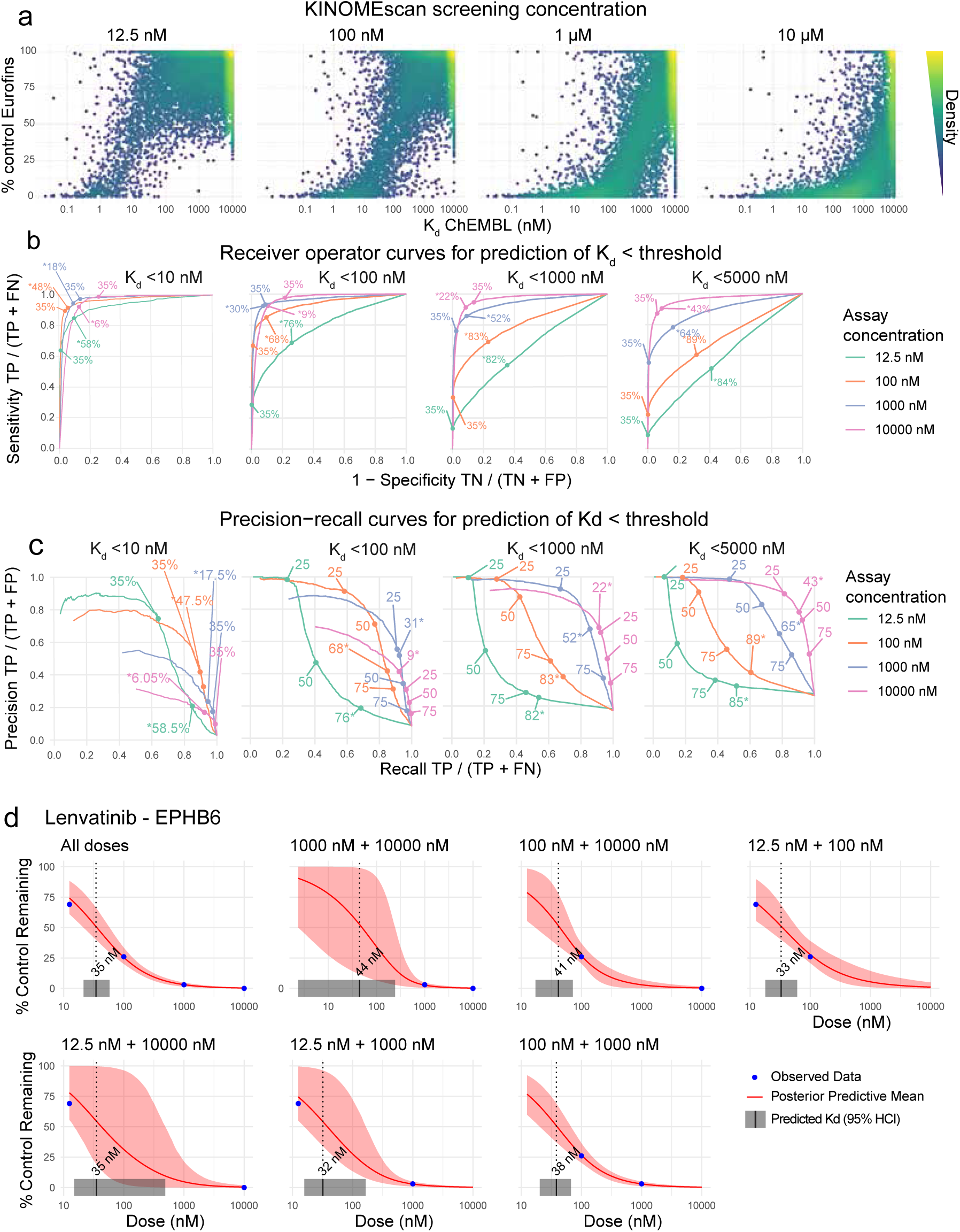
**(a)** Scatterplots showing % control values for the screening concentrations indicated with respect to K_d_ data in ChEMBL. **(b)** False positive and negative rates for all percent control thresholds are plotted (taking ChEMBL data as the ground truth) on receiver operator curves, and the % control corresponding to the point closest to the upper left corner of the plot is identified as the optimal threshold. The optimal % control thresholds (denoted by * and bold font) and 35% are shown for each curve. **(c)** Precision-recall curves for predicting the K_d_ values shown using screening data collected at each assay concentration. The optimal % control thresholds (denoted by * and bold font) and 35% are shown for each curve. **(d)** Dose-response fits from the Bayesian model for lenvatinib binding EPHB6 across all four doses (first plot) and all two dose combinations. The credible 95% interval for the fit is shaded in red and for the ^KS^K_d_ estimate in grey on the x-axis.

## References

1. Moret, N. et al. A resource for exploring the understudied human kinome for research and therapeutic opportunities. 2020.04.02.022277 Preprint at 10.1101/2020.04.02.022277 (2021).

2. Inhibitors Approved for Clinical Use | MRC PPU. https://www.ppu.mrc.ac.uk/list-clinically-approved-kinase-inhibitors.

3. Carles, F., Bourg, S., Meyer, C. & Bonnet, P. PKIDB: A Curated, Annotated and Updated Database of Protein Kinase Inhibitors in Clinical Trials. Molecules 23, 908 (2018).

4. Bournez, C. et al. Comparative Assessment of Protein Kinase Inhibitors in Public Databases and in PKIDB. Molecules 25, 3226 (2020).

5. Mullard, A. FDA approves 100th small-molecule kinase inhibitor. Nature Reviews Drug Discovery 24, 891–895 (2025).

6. Klaeger, S. et al. The target landscape of clinical kinase drugs. Science 358, (2017).

7. Davis, M. I. et al. Comprehensive analysis of kinase inhibitor selectivity. Nat Biotechnol 29, 1046–1051 (2011).

8. Anastassiadis, T., Deacon, S. W., Devarajan, K., Ma, H. & Peterson, J. R. Comprehensive assay of kinase catalytic activity reveals features of kinase inhibitor selectivity. Nat Biotechnol 29, 1039–1045 (2011).

9. Elkins, J. M. et al. Comprehensive characterization of the Published Kinase Inhibitor Set. Nat Biotechnol 34, 95–103 (2016).

10. Reinecke, M. et al. Chemical proteomics reveals the target landscape of 1,000 kinase inhibitors. Nat Chem Biol 1–9 (2023) doi:10.1038/s41589-023-01459-3.

11. Moret, N. et al. Cheminformatics Tools for Analyzing and Designing Optimized Small-Molecule Collections and Libraries. Cell Chemical Biology 26, 765–777.e3 (2019).

12. Zdrazil, B. et al. The ChEMBL Database in 2023: a drug discovery platform spanning multiple bioactivity data types and time periods. Nucleic Acids Research 52, D1180–D1192 (2024).

13. Nguyen, D.-T., et al. Pharos: Collating protein information to shed light on the druggable genome Nucleic Acids Research 45, D995–D1002 (2017).

14. Gilson, M. K. et al. BindingDB in 2015: A public database for medicinal chemistry, computational chemistry and systems pharmacology. Nucleic Acids Research 44, D1045–D1053 (2016).

15. Fabian, M. A. et al. A small molecule–kinase interaction map for clinical kinase inhibitors. Nat Biotechnol 23, 329–336 (2005).

16. Drewry, D. H. et al. Progress towards a public chemogenomic set for protein kinases and a call for contributions. PLOS ONE 12, e0181585 (2017).

17. Wells, C. I. et al. The Kinase Chemogenomic Set (KCGS): An Open Science Resource for Kinase Vulnerability Identification. Int J Mol Sci 22, E566 (2021).

18. Karaman, M. W. et al. A quantitative analysis of kinase inhibitor selectivity. Nat Biotechnol 26, 127–132 (2008).

19. Lahiry, P., Torkamani, A., Schork, N. J. & Hegele, R. A. Kinase mutations in human disease: interpreting genotype–phenotype relationships. Nat Rev Genet 11, 60–74 (2010).

20. Bikker, J. A., Brooijmans, N., Wissner, A. & Mansour, T. S. Kinase Domain Mutations in Cancer: Implications for Small Molecule Drug Design Strategies. J. Med. Chem. 52, 1493–1509 (2009).

21. Cheng, A. C., Eksterowicz, J., Geuns-Meyer, S. & Sun, Y. Analysis of kinase inhibitor selectivity using a thermodynamics-based partition index. J Med Chem 53, 4502–4510 (2010).

22. Bosc, N., Meyer, C. & Bonnet, P. The use of novel selectivity metrics in kinase research. BMC Bioinformatics 18, 17 (2017).

23. Sanfelice, D. et al. The Chemical Probes Portal – 2024: update on this public resource to support best-practice selection and use of small molecules in biomedical research. Nucleic Acids Res 53, D1663–D1669 (2025).

24. Essegian, D., Khurana, R., Stathias, V. & Schürer, S. C. The Clinical Kinase Index: A Method to Prioritize Understudied Kinases as Drug Targets for the Treatment of Cancer. Cell Rep Med 1, 100128 (2020).

25. Raschi, E., Vasina, V., Poluzzi, E. & De Ponti, F. The hERG K+ channel: target and antitarget strategies in drug development. Pharmacol Res 57, 181–195 (2008).

26. Dar, A. C., Das, T. K., Shokat, K. M. & Cagan, R. L. Chemical genetic discovery of targets and anti-targets for cancer polypharmacology. Nature 486, 80–84 (2012).

27. Yang, X. et al. Kinase Inhibition-Related Adverse Events Predicted from in vitro Kinome and Clinical Trial Data. J Biomed Inform 43, 376–384 (2010).

28. Roche. Alcensa Product Information. (2026).

29. Burwinkel, B., Shiomi, S., Al Zaben, A. & Kilimann, M. W. Liver glycogenosis due to phosphorylase kinase deficiency: PHKG2 gene structure and mutations associated with cirrhosis. Hum Mol Genet 7, 149–154 (1998).

30. Amemiya, T. et al. Elucidation of the molecular mechanisms underlying adverse reactions associated with a kinase inhibitor using systems toxicology. NPJ Syst Biol Appl 1, 15005 (2015).

31. Alectinib. in LiverTox: Clinical and Research Information on Drug-Induced Liver Injury (National Institute of Diabetes and Digestive and Kidney Diseases, Bethesda (MD), 2012).

32. Dark Kinase Knowledgebase: an online compendium of knowledge and experimental results of understudied kinases | Nucleic Acids Research | Oxford Academic. https://academic.oup.com/nar/article/49/D1/D529/5932842?login=false.

33. Sharma, K. R., Colvis, C. M., Rodgers, G. P. & Sheeley, D. M. Illuminating the druggable genome: Pathways to progress. Drug Discov Today 29, 103805 (2024).

34. Vijayan, R. S. K. et al. Conformational Analysis of the DFG-Out Kinase Motif and Biochemical Profiling of Structurally Validated Type II Inhibitors. J. Med. Chem. 58, 466–479 (2015).

35. Zhang, J., Yang, P. L. & Gray, N. S. Targeting cancer with small molecule kinase inhibitors. Nat Rev Cancer 9, 28–39 (2009).

36. Berman, H. M. et al. The Protein Data Bank. Nucleic Acids Res 28, 235–242 (2000).

37. Hanson, S. M. et al. What makes a kinase promiscuous for inhibitors? Cell Chem Biol 26, 390–399.e5 (2019).

38. Manning, G., Whyte, D. B., Martinez, R., Hunter, T. & Sudarsanam, S. The Protein Kinase Complement of the Human Genome. Science 298, 1912–1934 (2002).

39. Wang, X., Ding, J. & Meng, L. PI3K isoform-selective inhibitors: next-generation targeted cancer therapies. Acta Pharmacol Sin 36, 1170–1176 (2015).

40. Lindberg, M. F. & Meijer, L. Dual-Specificity, Tyrosine Phosphorylation-Regulated Kinases (DYRKs) and cdc2-Like Kinases (CLKs) in Human Disease, an Overview. Int J Mol Sci 22, 6047 (2021).

41. Song, M. et al. Cdc2-like kinases: structure, biological function, and therapeutic targets for diseases. Sig Transduct Target Ther 8, 1–25 (2023).

42. Schröder, M. et al. DFG-1 Residue Controls Inhibitor Binding Mode and Affinity, Providing a Basis for Rational Design of Kinase Inhibitor Selectivity. J. Med. Chem. 63, 10224–10234 (2020).

43. Rodriguez, S. et al. Genome-encoded cytoplasmic double-stranded RNAs, found in C9ORF72 ALS-FTD brain, propagate neuronal loss. Sci Transl Med 13, eaaz4699 (2021).

44. Konig, L.E. et al. TYK2 as a novel therapeutic target in Alzheimer’s Disease with TDP-43 inclusions. Nature Neuroscience (Under review.).

45. Wong, P.-M., Puente, C., Ganley, I. G. & Jiang, X. The ULK1 complex. Autophagy 9, 124–137 (2013).

46. Levin-Salomon, V., Bialik, S. & Kimchi, A. DAP-kinase and autophagy. Apoptosis 19, 346–356 (2014).

47. Selvaraj, V., et al. Baricitinib in hospitalised patients with COVID-19: A meta-analysis of randomised controlled trials. eClinicalMedicine 49, (2022).

48. Zhang, Z., Yang, X., Song, Y.-Q. & Tu, J. Autophagy in Alzheimer’s disease pathogenesis: Therapeutic potential and future perspectives. Ageing Research Reviews 72, 101464 (2021).

49. Jung, C. H., Ro, S.-H., Cao, J., Otto, N. M. & Kim, D.-H. mTOR regulation of autophagy. FEBS Lett 584, 1287–1295 (2010)

50. Sakamaki, J. et al. Bromodomain Protein BRD4 Is a Transcriptional Repressor of Autophagy and Lysosomal Function. Mol Cell 66, 517–532.e9 (2017).

51. Petrova, V., et al. Multi-Kinase Inhibition of STE20 Kinases is Neuroprotective Against Chemotherapy-Induced Axon Degeneration. SSRN Scholarly Paper at 10.2139/ssrn.5380584 (2025).

52. Bos, P. H. et al. Development of MAP4 Kinase Inhibitors as Motor Neuron-Protecting Agents. Cell Chem Biol 26, 1703–1715.e37 (2019).

53. Davis, A., Tinker, A. V. & Friedlander, M. ‘Platinum resistant’ ovarian cancer: what is it, who to treat and how to measure benefit? Gynecol Oncol 133, 624–631 (2014).

54. Granados, M. L., Hudson, L. G. & Samudio-Ruiz, S. L. Contributions of the Epidermal Growth Factor Receptor to Acquisition of Platinum Resistance in Ovarian Cancer Cells. PLoS One 10, e0136893 (2015).

55. Zhao, J. et al. FGFR3 phosphorylates EGFR to promote cisplatin-resistance in ovarian cancer. Biochemical Pharmacology 190, 114536 (2021).

56. Wilken, J. A. et al. EGFR/HER-targeted therapeutics in ovarian cancer. Future Med Chem 4, 447–469 (2012).

57. Zeng, J. et al. Protein kinases PknA and PknB independently and coordinately regulate essential Mycobacterium tuberculosis physiologies and antimicrobial susceptibility. PLOS Pathogens 16, e1008452 (2020).

58. Thongdee, P. et al. Virtual Screening Identifies Novel and Potent Inhibitors of Mycobacterium tuberculosis PknB with Antibacterial Activity. J. Chem. Inf. Model. 62, 6508–6518 (2022).

59. Xu, J. et al. A novel protein kinase inhibitor IMB-YH-8 with anti-tuberculosis activity. Sci Rep 7, 5093 (2017).

60. Larkins-Ford, J. et al. Systematic measurement of combination-drug landscapes to predict *in vivo* treatment outcomes for tuberculosis. Cell Systems 12, 1046–1063.e7 (2021).

61. Singer, J. W. et al. Comprehensive kinase profile of pacritinib, a nonmyelosuppressive Janus kinase 2 inhibitor. J Exp Pharmacol 8, 11–19 (2016).

62. Keenan, S. M. & Welsh, W. J. Characteristics of the Plasmodium falciparum PK5 ATP-binding site: implications for the design of novel antimalarial agents. J Mol Graph Model 22, 241–247 (2004).

63. Ou, S.-H. I., Bartlett, C. H., Mino-Kenudson, M., Cui, J. & Iafrate, A. J. Crizotinib for the Treatment of ALK-Rearranged Non-Small Cell Lung Cancer: A Success Story to Usher in the Second Decade of Molecular Targeted Therapy in Oncology. Oncologist 17, 1351–1375 (2012).

64. Shaw, A. T. et al. Crizotinib in ROS1-rearranged advanced non-small-cell lung cancer (NSCLC): updated results, including overall survival, from PROFILE 1001. Ann Oncol 30, 1121–1126 (2019).

65. Hafner, M. et al. Multiomics Profiling Establishes the Polypharmacology of FDA-Approved CDK4/6 Inhibitors and the Potential for Differential Clinical Activity. Cell Chem Biol 26, 1067–1080.e8 (2019).

66. Dar, A. C., Das, T. K., Shokat, K. M. & Cagan, R. L. Chemical genetic discovery of targets and anti-targets for cancer polypharmacology. Nature 486, 80–84 (2012).

67. Zhao, L., Zhao, J., Zhong, K., Tong, A. & Jia, D. Targeted protein degradation: mechanisms, strategies and application. Sig Transduct Target Ther 7, 113 (2022).

68. Wodicka, L. M. et al. Activation State-Dependent Binding of Small Molecule Kinase Inhibitors: Structural Insights from Biochemistry. Chemistry & Biology 17, 1241–1249 (2010).

69. Abril-Pla, O. et al. PyMC: a modern, and comprehensive probabilistic programming framework in Python. PeerJ Comput. Sci. 9, e1516 (2023).

70. Masters, J. R. et al. Short tandem repeat profiling provides an international reference standard for human cell lines. Proceedings of the National Academy of Sciences 98, 8012–8017 (2001).

71. Mills, C. E. et al. Multiplexed and reproducible high content screening of live and fixed cells using Dye Drop. Nat Commun 13, 6918 (2022).

72. Hafner, M., Niepel, M., Chung, M. & Sorger, P. K. Growth rate inhibition metrics correct for confounders in measuring sensitivity to cancer drugs. Nature methods 13, 521–7 (2016).

73. Abraham, M. J. et al. GROMACS: High performance molecular simulations through multi-level parallelism from laptops to supercomputers. *SoftwareX* 1–2, 19–25 (2015).

74. Gromacs: A parallel computer for molecular dynamics simulations – ScienceOpen. https://www.scienceopen.com/book?vid=59290415-039c-4900-8e95-d649687a2473.

75. Guvench, O. et al. CHARMM Additive All-Atom Force Field for Carbohydrate Derivatives and Its Utility in Polysaccharide and Carbohydrate–Protein Modeling. J. Chem. Theory Comput. 7, 3162–3180 (2011).

76. Malde, A. K. et al. An Automated Force Field Topology Builder (ATB) and Repository: Version 1.0. J. Chem. Theory Comput. 7, 4026–4037 (2011).

77. Haug, E. J., Arora, J. S. & Matsui, K. A steepest-descent method for optimization of mechanical systems. J Optim Theory Appl 19, 401–424 (1976).

78. Parrinello, M. & Rahman, A. Polymorphic transitions in single crystals: A new molecular dynamics method. Journal of Applied Physics 52, 7182–7190 (1981).

79. Gunsteren, W. F. V. & Berendsen, H. J. C. A Leap-frog Algorithm for Stochastic Dynamics. Molecular Simulation https://doi.org/10.1080/08927028808080941 (1988) doi:10.1080/08927028808080941.

80. Darden, T., York, D. & Pedersen, L. Particle mesh Ewald: An N⋅log(N) method for Ewald sums in large systems. The Journal of Chemical Physics 98, 10089–10092 (1993).

